# TGFβR Inhibition Represses TGF-β1 Initiated Keratin-7 Expression in Human Salivary Gland Progenitor Cells

**DOI:** 10.1101/2021.10.08.463706

**Authors:** Eric W. Fowler, Emmett V. Venrooy, Robert L. Witt, Xinqiao Jia

## Abstract

Towards the goal of engineering an implantable salivary gland for the treatment of xerostomia, we culture primary human salivary gland stem/progenitor cells (hS/PCs) in hyaluronic acid (HA)-based hydrogels containing a covalently conjugate integrin-binding peptide (RGDSP). We characterize how RGDSP affects hS/PC phenotype and discover the presence of cells expressing both amylase and keratin-7 (K7) in our 3D cultures. Typically, amylase is expressed by acinar cells, and K7 is found in ducts. After assaying an array of transforming growth factor-β (TGF-β) superfamily members, we find increased expression of TGF-β1 and growth/differentiation factor-15 (GDF-15) in RGDSP cultures. However, 2D model studies confirm that only TGF-β1 is required to induce K7 expression in hS/PCs. We then demonstrate that with pharmacological inhibition of TGF-β signaling, K7 expression is repressed while amylase expression is maintained in RGDSP cultures. Thus, TGF-β signaling regulates K7 expression in hS/PCs, and modulation of TGF-β signaling is essential for the regeneration of salivary gland function.

## 1. Introduction

More than 600,000 head and neck cancer cases are diagnosed yearly, for which radiation therapy (RT) is a common successful therapeutic.^[1]^ Unfortunately, the salivary gland (SG) can be damaged in the process of RT due to its proximity to the field of irradiation.^[1a, 2]^ SGs are composed of acinar cells that produce saliva and ductal cells that modify the saliva and direct its flow.^[1a, 3]^ When acinar cells are damaged, they reduce or cease saliva production, which results in SG dysfunction that impedes one’s ability to speak, swallow, masticate, and compromises oral health, ultimately impairing patients’ well-being.^[4]^Currently, approved therapies aim to increase saliva production from the remaining acinar cells; however, these methods do not offer regenerative stimuli to increase the acinar cell population.^[5]^ There is a critical need for treatments that can restore SG function and improve patients’ quality of life.

Tissue engineering offers regenerative opportunities for the treatment of SG dysfunction; however, the isolation and *in vitro* expansion of saliva producing acinar cells has remained challenging. For example, *in vitro* culture of SG cells has been reported to lead to loss of the secretory acinar identity.^[6]^ Recently, it was shown that stress or injury can activate the phenotypic plasticity of acinar cells, stimulating their expression of ductal specific keratins, such as keratin-7 (K7) and keratin-19 (K19).^[6a]^ The signaling pathways behind acinar cell plasticity observed during SG culture are not well understood. However, previous research has implicated dysregulation of Rho-associated protein kinase (ROCK),^[6e]^ epidermal growth factor receptor (EGFR),^[7]^ proto-oncogene c-Src (Src) and p38 mitogen-activated protein kinase (MAPK),^[6b-d]^ or transforming growth factor-beta (TGF-β) signaling to be involved in the increased ductal identity in primary SG cultures.^[1a, 8]^

TGF-β signaling is of interest as TGF-β superfamily members are reported to regulate K7 expression in various contexts.^[9]^ In addition, TGF-β induces cell-cycle arrest in most non-malignant epithelial cells.^[10]^ Cells undergoing irreversible cell-cycle arrest, or senescence, often secrete high levels of inflammatory cytokines, proteases, protease inhibitors, and extracellular matrix (ECM) proteins including: interleukin 6 (IL-6), interleukin-8 (IL-8), matrix metalloproteinase-1 (MMP-1), plasminogen activator inhibitor-1 (PAI-1), and fibronectin, the senescence-associated secretory phenotype (SASP).^[10a, 11]^ Although the secreted factors of SASP are highly dynamic and varied across tissues, a recent proteomic study identified growth/differentiation factor-15 (GDF-15), a divergent member of the TGF-β superfamily expressed in response to cellular stress, as a predictive biomarker maintained across SASPs.^[11b]^ Notably, elevated GDF-15 expression has been reported in irradiated SGs,^[12]^ where senescence is known to play a major role in disrupting SG regeneration.^[13]^ Notwithstanding, elevated TGF-β expression has also been found in irradiated SGs, and TGF-β signaling is reported to be a main effector of SASP.^[11a, 14]^

TGF-β further plays a dichotomous role in driving disease progression, initially acting as a cytostatic tumor suppressor and later switching to a tumorigenic driver by inducing cell motility and epithelial-to-mesenchymal transition (EMT).^[10b, 15]^ EMT is an extreme state of phenotypic plasticity, where epithelial cells de/transdifferentiate and take on the characteristics of motile mesenchymal cells.^[13c, 15-16]^ Transient EMT is often exhibited during tissue repair, but sustained or repeated states of EMT can produce stiff fibrotic tissue accumulation, or fibrosis.^[13c, 17]^ As the central mediator of fibrosis, canonical TGF-β signaling is carried out through the phosphorylation and nuclear import of SMAD 2 and SMAD 3 (SMAD 2/3).^[15-16, 18]^ Interestingly, SMAD 2/3 signaling can also be influenced by the local ECM microenvironment through potentiating interactions with the mechanically activated Yes Associated Protein (YAP).^[18-19]^ There is rising evidence that YAP contributes to fibrotic diseases by forming a feedback loop by driving the expression of pro-fibrotic cytokines and nuclear retention of SMAD 2/3, which ultimately results in a stiff microenvironment that further potentiates YAP signaling.^[19a, 20]^

We have previously reported hyaluronic acid (HA)-based 3D matrices that support the 3D culture of primary human salivary stem progenitor cell (hS/PC) under defined conditions and identified a mechanical stiffness regime that promotes the development of multicellular spheroids.^[21]^ ECM composition is a principal regulator of SG development and differentiation.^[7, 22]^ Recently, we demonstrated that incorporating the fibronectin-derived RGDSP integrin ligand in HA matrices promoted the rapid proliferation of amylase expressing hS/PCs.^[23]^ Providing integrin-mediated adhesion could be expected to increase nuclear YAP, by enhancing mechanotransduction, to promote TGF-β signaling in RGDSP cultures.^[19a, 24]^

Here, we investigate how RGDSP directs hS/PC phenotype and report on the contributions from SMAD2/3 and YAP signaling. We show that TGF-β1 can induce K7 expression in hS/PCs and characterize how YAP signaling participates in both TGF-β signaling and GDF-15 expression. We find that K7 expression arises under conditions that are characteristic of SASP and EMT. After providing a mechanism for high GDF-15 expression observed in RGDSP cultures, we demonstrate that K7 expression can be repressed in both 2D and 3D models by TGFβR inhibition. K7 is widely utilized as a terminal ductal SG marker, yet K7 expression could indicate a state of cellular stress experienced during primary SG culture. This work provides context to the K7 expressing phenotype and demonstrates how to mitigate K7 expression.

## 2. Results

### 2.1. RGDSP promotes rapid development of multicellular epithelial structures

Salivary gland development and differentiation is instructed by both matrix and cell-derived interactions.^[22a, 22c]^ To this end, we performed super-resolution microscopy to characterize the temporal dynamics of cell-matrix and cell-cell interactions arising in the HA and RGDSP cultures over time (**Figure** 1a). When encapsulated as single cells, hS/PCs formed multicellular spheroids over 14 days. The expression of keratin-5 (K5), a SG progenitor marker,^[4b, 25]^ was maintained in both culture conditions and localized basally as cell division proceeded. Compared to HA controls, RGDSP hydrogels promoted the development of larger hS/PC structures as early as day 3 (Figure 1a-b) before significant proliferation was detected (Figure S2a). The structures continued to grow, and on day 14, spheroids in RGDSP cultures averaged twice the size of those found in HA cultures (Figure 1b). In both HA and RGDSP cultures, circularity increased as spheroids developed in size, and RGDSP cultures maintained higher circularity from days 3-14 (Figure 1c). The F-actin derived circularity was largely influenced by the filopodia extending from the hS/PC structures at early time points. On day 1, hS/PCs were found predominantly in the single-cell state and extended filopodia ~600 nm in length in both HA and RGDSP cultures. By day 3, extended filopodia were only maintained in HA hydrogels, and a significant decrease in filopodia length was observed in RGDSP cultures (Figure 1d-e).

**Figure 1.**
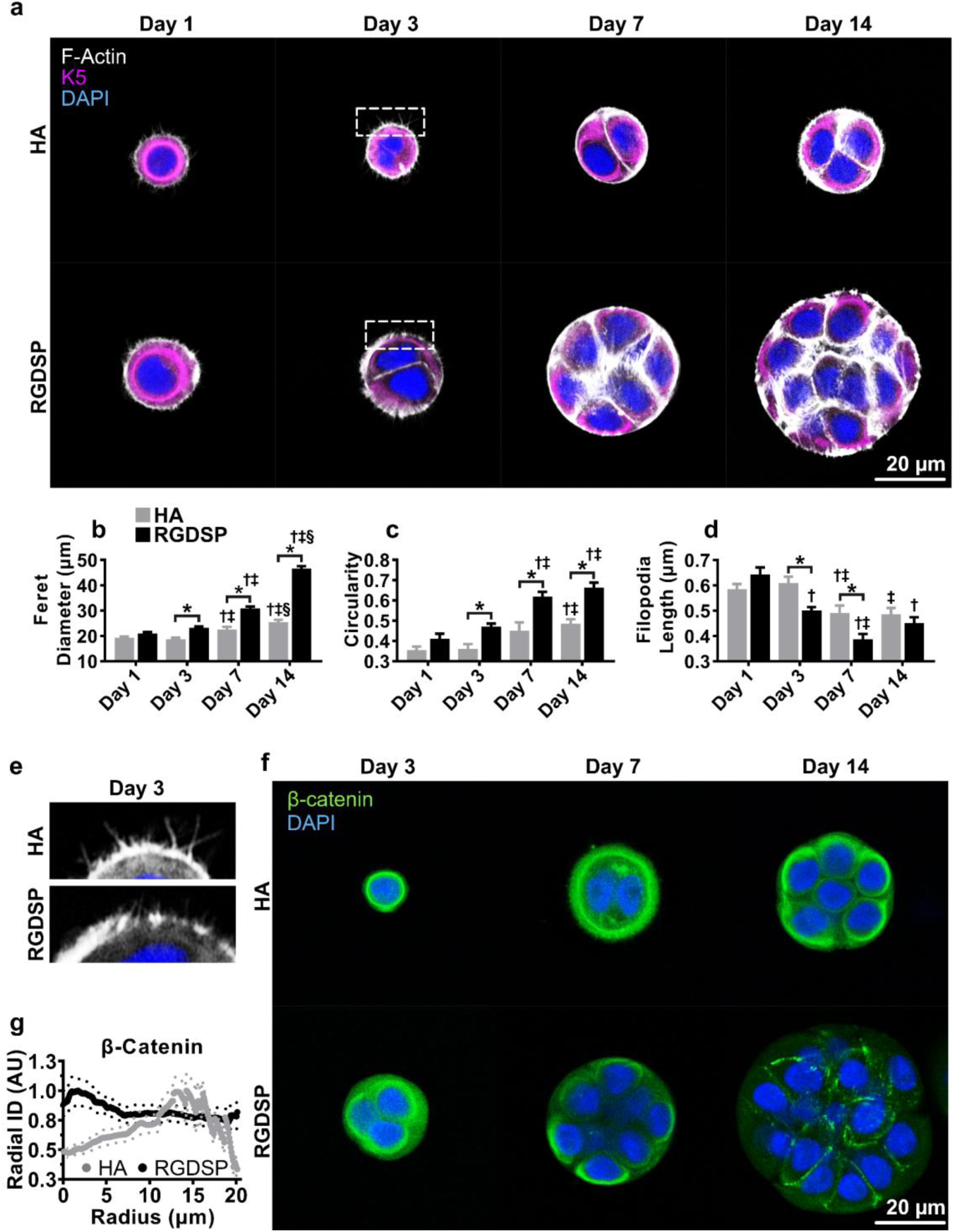
Hydrogel microenvironment drives temporal cell-cell and cell-ECM dynamics. **(a)** Fluorescent microscopy indicating localization of K5 and F-actin on days 1, 3, 7, and 14 of culture. **(b-d)** Morphological analysis of hS/PC ferret diameter (b), circularity (c), and filopodia length (d) was performed from fluorescent microscopy. **(e)** Enlargement of day 3 filopodia. **(f)** β-catenin localization at day 3, day 7, and day 14. **(g)** Radial profile detailing β-catenin localization at day 14. Error bars represent SEM in all cases. Two-way ANOVAs were performed on (c-e) data sets, followed by Tukey’s multiple comparison test. ^*^ indicates *p* < 0.05 between HA and RGDSP at the same time point. ^†, ‡, §^ indicates *p <* 0.05 from day 1, 3, or day 7 measurements of the same data set, respectively.

The increased circularity of spheroids formed in RGDSP cultures suggested enhanced cell-cell adhesion. The predominant way in which cell-cell adhesion is realized is through the cadherin/catenin adhesion complexes.^[26]^ In HA cultures, β-catenin, which localizes to epithelial adherins junctions,^[26a, 26b]^ was expressed in a diffuse pattern throughout the cytoplasm, irrespective of the stage of spheroid development (Figure 1f-g). β-catenin presented in RGDSP cultures with a similarly diffuse pattern until day 14, where it was highly localized to the apical and lateral membranes. Thus, both HA and RGDSP matrices promoted filopodia extension into the surrounding matrix, but only RGDSP cultures formed defined adherins junctions, indicating enhanced cell-cell adhesion.

### 2.2. RGDSP cultures developed a mixed phenotype with increased K7 and α-amylase expression

To assess whether the presented microenvironmental cues promoted a pro-ductal or pro-acinar phenotype, we next characterized the temporal gene expression dynamics of HA and RGDSP cultures with qPCR (**Figure** 2a).

**Figure 2.**
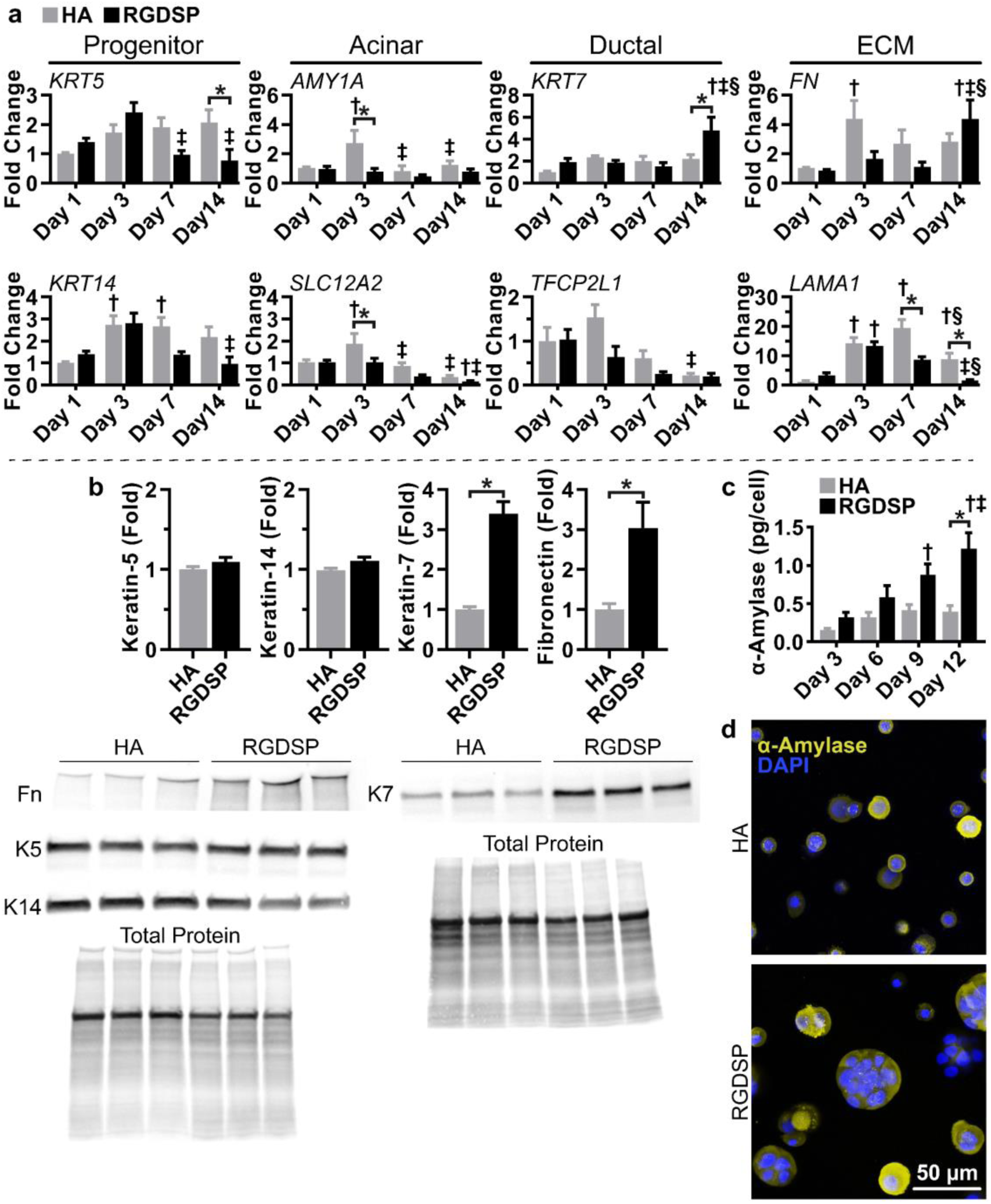
RGDSP promotes a mixed lineage with increased K7 and α-amylase expression. **(a)** Temporal gene expression profile via RT-qPCR analysis of *KRT5, KRT14, SLC12A2, KRT7, TFCP2L1, FN*, and *LAMA1* at day 1, 3, 7, 14 of culture. **(b)** Western blot analysis was conducted on day 14 for K5, K14, K7, and fibronectin. **(c)** Secreted α-amylase quantified on days 3, 6, 9, and 12 by ELISA. Triplicate measurements are presented from technical replicates extracted from different hydrogels. **(d)** Immunocytochemistry (ICC) detailing uniform expression of α-amylase throughout HA and RGDSP spheroids at day 14 of culture. Error bars represent SEM in all cases. Two-way ANOVAs were conducted on data presented in (a) and (c), followed by Tukey’s multiple comparison test. ^*^ indicates *p* < 0.05 between HA and RGDSP at the same time points. ^†, ‡, §^ indicates *p* < 0.05 from day 1, 3, and day 7 measurements of the same data set, respectively for (a) and days 3, 6, 9 for (c). Two-tailed Student’s t-tests were conducted on data appearing in (b), where ^*^ indicates *p* < 0.05.

Keratins expressed by SG progenitors, K5 (K*RT5*) and keratin-14 (*KRT14*),^[4b, 25]^ presented similar temporal profiles that were unique to each ECM condition. There was a moderate increase in *KRT5*/*KRT14* expression in HA cultures following day 1 that was sustained to day 14. *KRT5*/*KRT14* expression in RGDSP cultures similarly increased on day 3; however, *KRT5/KRT14* expression decreased thereafter, with *KRT5* expression levels decreasing significantly below expression observed in HA cultures. Analysis of the gene expression profile of acinar markers α-amylase (*AMY1A*) and the Na-K-Cl ion transporter (*SLC12A2*) revealed separate trends.^[1a, 4b, 25a]^ *AMY1A* expression was relatively stable across time in both HA and RGDSP cultures. Although in HA cultures, *AMY1A* expression was elevated on day 3, it returned to basal expression levels on day 7. HA cultures maintained *SLC12A2* expression through day 7; however, on day 14, *SLC12A2* was downregulated with respect to its expression on day 1. Comparatively, loss of *SLC12A2* expression occurred earlier in RGDSP cultures (day 7), with a significant reduction in expression occurring on day 7 that was significantly below the expression observed in HA cultures at day 7, and decreased further on day 14. The ductal marker keratin-7 (*KRT7*) and a transcription factor involved in duct regulation (*TFCP2L1*),^[1a, 4b, 27]^ like the expression of acinar markers, were not expressed in a phenotype-specific manner. HA and RGDSP cultures stably expressed *KRT7* until day 14, where RGDSP promoted a 2.2 fold higher *KRT7* expression relative to the day 14 HA cultures (4.8 fold increased when normalized to the expression of HA cultures on day 1). However, a corresponding increase in the expression of the K7 dimerization partners, keratin-18 (*KRT18*) and keratin-19 (*KRT19*),^[28]^was not detected in RGDSP cultures on day 14 (Figure S2b). *TFCP2L1* expression was maintained in HA cultures until day 14, where it was downregulated relative to the normalized values of HA cultures on day 1. RGDSP cultures initiated the downregulation of *TFCP2L1* at day 3, and it was further decreased, relative to RGDSP on day 1, through days 7-14; however, only days 3-7 were significantly different between HA and RGDSP cultures, due to a similar loss of expression occurring in HA cultures at day 14. The gene expression dynamics of ECM proteins which are important in SG development and differentiation were also examined, including fibronectin (*FN*), required in SG development,^[22a]^ and laminin, alpha-1 subunit (*LAMA1*), which is associated with the maintenance of the acinar phenotype.^[22b, 22d]^ *FN* and *LAMA1* were upregulated at days 3-7 under both culture conditions relative to the expression level by HA cultures on day 1; however, by day 14, *FN* was upregulated, and *LAMA1* was downregulated in the RGDSP cultures.

Protein level confirmation of these findings was made by western blot analyses of K5, K14, K7, and fibronectin (Figure 2b). Though *KRT5* mRNA expression was significantly lower in RGDSP cultures on day 14, we did not detect a loss in K5 expression on day 14 at the protein level. Furthermore, the expression of K14, the predominant heterodimeric partner of K5,^[28]^ was also retained in the RGDSP cultures at day 14. However, K7 and fibronectin were increased by ~3-fold (*p* < 0.05) in RGDSP cultures, in agreement with the corresponding mRNA expression profiles. As increased ductal K7 expression might be associated with the loss of the secretory amylase protein, amylase expression was evaluated via ELISA. RGDSP cultures produced approximately 2-fold more amylase per cell on day 12 than HA cultures (Figure 2c), suggesting a mixed population of pro-ductal and pro-acinar spheroids. However, the immunostaining pattern of α-amylase indicated that α-amylase was uniformly expressed throughout both HA and RGDSP cultures and was not restricted to a specific cell population (Figure 2d). Collectively, these findings indicate that RGDSP increased secretory amylase expression while also enhancing expression of the ductal marker K7.

### 2.3. TGF-β1 expression and nuclear SMAD 2/3 localization correspond to increased K7 expression

To promote a secretory acinar phenotype and repress the development of the undesirable K7 phenotype in RGDSP cultures, we set out to identify potential upstream regulators of K7. Literature mining indicated that members of the TGF-β family are responsible for activating K7 expression in other contexts.^[9, 29]^ We used an immunoblot array to evaluate the production of TGF-β superfamily members in the medium collected from HA and RGDSP cultures on day 6. Increased expression of TGF-β1 and GDF-15, relative to HA cultures, was detected in the medium of RGDSP cultures (Figure S2c). We then conducted TGF-β1 and GDF-15 ELISAs to confirm the array findings and assess the temporal cytokine expression dynamics (**Figure** 3a).

**Figure 3.**
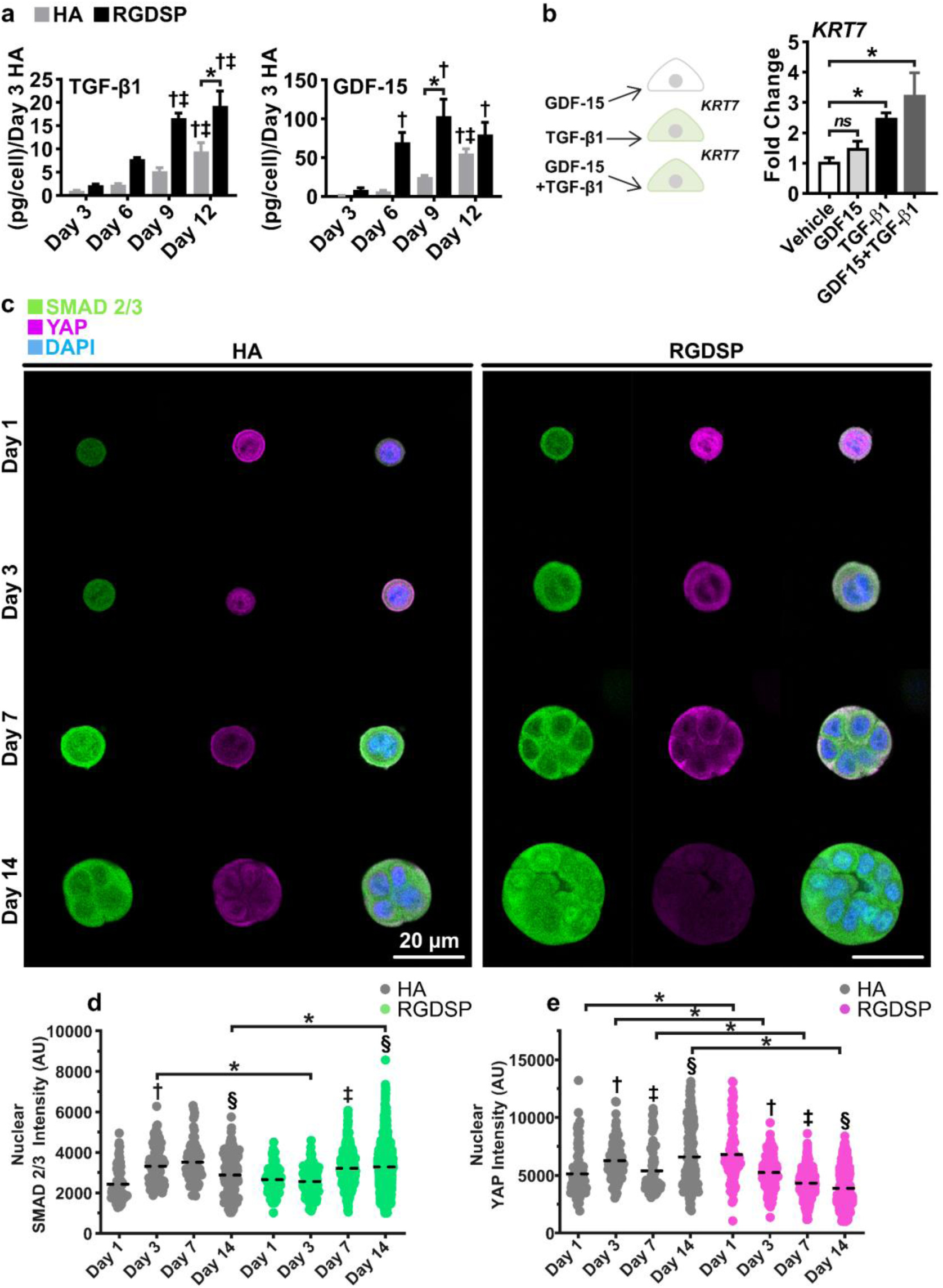
RGDSP cultures expressed high levels of TGF-β1 and GDF-15 that correlate with increased nuclear SMAD 2/3. Only TGF-β1 is required for *KRT7* expression in 2D cultures. **(a)** ELISA analyses were conducted targeting soluble TGF-β1 and GDF-15 in cell culture medium after 3, 6, 9, and 12 days of culture. **(b)** hS/PCs were cultured with TGF-β1, GDF-15, and TGF-β1+GDF-15 for 48 h on 2D substrates, and the expression of K7 was resolved with qPCR. Significance determined from one-way ANOVA followed by a Dunnett’s test, ^*^ indicates *p* < 0.05 relative to vehicle control. **(c)** ICC detailing hS/PC nuclear YAP and SMAD 2/3 on days 1, 3, 7, and 14 of culture in HA and RGDSP hydrogels. Volumetric fluorescent microscopy was conducted, and representative single plane images are presented. **(d-e)** 3D rendering was performed with Imaris 3D-4D software for the quantification of nuclear SMAD 2/3 (d) and YAP (e) from HA and RGDSP cultures on days 1, 3, 7, and 14 of culture. Filled circles represent individual nuclei, and the dashed black line indicates the mean value of each data set. Error bars represent SEM in all cases. Two-way ANOVAs were performed on data presented in (a) and (d-e) followed by Tukey’s multiple comparison test. ^*^ indicates *p* < 0.05 between HA and RGDSP at the same time points. For (a), ^†, ‡, §^ indicates *p* < 0.05 from day 3, 6, and day 9 measurements of the same data set, respectively. In (d-e), ^†, ‡, §^ indicates *p* < 0.05 from day 1, 3, and day 7 measurements of the same data set, respectively.

From days 3-12, HA and RGDSP cultures increased TGF-β1 expression by 9- and 19-fold, respectively. However, RGDSP cultures maintained at least a 2-fold higher level of TGF-β1 expression relative to HA cultures throughout the entire culture period. The difference in GDF-15 expression between HA and RGDSP cultures was more dramatic. A 7- and a 25-fold increase in GDF-15 expression, relative to the initial day 3 HA levels, was observed at days 6 and 9 in HA cultures, but RGDSP cultures increased 70- to 103-fold over the same time period. However, GDF-15 levels continued to rise in HA cultures, and by day 12 were no longer expressed at significantly different levels in HA and RGDSP cultures.

Since TGF-β1 and GDF-15 were both highly expressed in RGDSP cultures alongside K7, we examined if TGF-β1 (10 ng mL^**-1**^) or GDF-15 (100 ng mL^**-1**^) was capable of inducing *KRT7* expression. 2D hS/PC cultures were treated with each cytokine independently and in combination for 48 h before assessing *KRT7* mRNA expression (Figure 3b). Treatment with GDF-15 alone did not alter *KRT7* transcript levels; however, TGF-β1 induced *KRT7* expression and sustained *KRT7* stimulation in the presence of GDF-15 (i.e., GDF-15+TGF-β1 treatment group). Thus, TGF-β1 is capable of stimulating *KRT7* expression and is differentially expressed in RGDSP cultures.

Based on the evidence that GDF-15 is highly predictive of a SASP,^[11b]^ we further considered an indirect role for TGF-β1 and GDF-15 to modulate *KRT7* expression by stimulating expression of SASP inflammatory factors. To this end, we compared the expression of a panel of SASP genes in HA and RGDSP cultures on day 14 to those stimulated by TGF-β1, GDF-15, and TGF-β1+GDF-15 in 2D cultures (Figure S3a-b). The expression of senescent and inflammatory factors was largely governed by TGF-β1 in 2D cultures, while the activity of GDF-15 was limited to increasing the expression of *IL1B*. In 3D cultures, RGDSP hydrogels stimulated high levels of the SASP-associated protease MMP-1 (Figure S3c).^[11b, 11c]^ In regards to the cell cycle inhibitors that are indicative of senescence, TGF-β1 did not stimulate cyclin-dependent kinase inhibitor 1A (*CDKN1A*, P21), yet cyclin-dependent kinase inhibitor 2A (*CDKN2A*, P16) and tumor protein P53 (*TP53*) were upregulated in 2D cultures.^[11, 13a, 13b]^ However, RGDSP cultures did not activate *CDKN1A, CDKN2A*, or *TP53* expression, suggesting growth arrest had not occurred. In 2D cultures, TGF-β1 activated *IL6* expression, yet RGDSP cultures increased *IL8* expression without stimulating *IL6* expression. Thus, *KRT7* expression was separately correlated with *IL6* and *IL8* expression but not maintained between 2D and 3D conditions, suggesting interleukins did not regulate *KRT7* expression. Moreover, elevated expression of the TGF-β1 target gene *SERPINE1* was observed in RGDSP cultures,^[9d, 16]^ further suggesting that TGF-β1 was the dominant factor governing emergence of the K7 phenotype.

Next, we characterized TGF-β signaling in RGDSP cultures. Furthermore, the interplay of TGF-β signaling and integrin-mediated mechanotransduction was investigated by targeting SMAD 2/3, the canonical TGF-β pathway signal transducers,^[15-16, 18]^ and YAP, a mechanically sensitive co-transcription factor,^[18-19]^ by conducting double immunofluorescence (Figure 3c). Quantification of nuclear SMAD 2/3 and YAP expression indicated distinct temporal dynamics between HA and RGDSP cultures (Figure 3d-e). HA cultures exhibited balanced temporal profiles of SMAD 2/3 and YAP expression, where enhanced nuclear SMAD 2/3 and YAP were observed at day 3. In HA cultures after day 3, nuclear SMAD 2/3 and YAP were inversely localized. YAP decreased at day 7 while SMAD 2/3 increased, and on day 14, nuclear YAP increased, yet nuclear SMAD 2/3 decreased. Conversely, in RGDSP cultures, SMAD 2/3 and YAP expression was highly dysregulated. RGDSP promoted peak nuclear YAP on day 1, whereafter it continuously decreased along with cytoplasmic YAP (Figure S4a). As nuclear YAP decreased in RGDSP cultures, nuclear SMAD 2/3 increased significantly at day 7 and furthermore at day 14, resulting in significantly higher levels at day 14 than observed in HA cultures. However, cytoplasmic SMAD 2/3 levels did not significantly increase in RGDSP cultures (Figure S4b). These findings confirm that the increased TGF-β signaling in RGDSP cultures corresponds to the rising TGF-β1 levels in the medium and increased K7 expression in RGDSP cultures. Conversely, RGDSP cultures do not sustain elevated YAP expression to drive TGF-β signaling and K7 expression.

The immunofluorescence determined nuclear SMAD 2/3 and YAP expression was confirmed by comparing their temporal localizations with the gene expression z-scores of the HA and RGDSP cultures, including TGF-β and YAP targets (Figure S4c). Irrespective of nuclear SMAD2/3 and YAP expression, HA and RGDSP cultures were defined by initial *ADAM10*/*ADAM17* activation with the expression of EGFR ligands, followed by a transition to *MMP1* expression as culture time proceeded. In HA cultures, the appearance of nuclear SMAD 2/3 and YAP together at day 3 dominated the transcriptional events where *KRT7, FN*, and *ITGAV* were maximally expressed. However, in RGDSP cultures, nuclear SMAD 2/3 and YAP were dysregulated, and nuclear SMAD 2/3 expression increased to day 14, corresponding to the highest *KRT7, FN*, and *ITGAV* expression. In RGDSP cultures, nuclear YAP expression clearly proceeded *YAP1* and *CTGF* expression; however, in HA cultures, the co-occupancy of nuclear SMAD2/3 and YAP together at day 3 maximized *YAP1* and *CTGF*. Thus, differences in nuclear SMAD2/3 and YAP expression were also seen at the gene level; however, the co-regulation of selected genes confirms the temporal localization observed with microscopy.

### 2.4. Loss of nuclear YAP stimulates GDF15 expression and downregulates TGF-β target genes

We have shown that TGF-β1 can stimulate K7 expression (Figure 2b) in hS/PCs, yet YAP is involved in maintaining the stem/progenitor status of various tissues, and loss of nuclear YAP expression could be sufficient to induce differentiation.^[30]^ To this end, a series of YAP inhibition studies were performed to investigate TGF-β dependent YAP signaling and how the loss of nuclear YAP could contribute to the observed K7 expressing phenotype development. 2D hS/PC cultures were treated with the YAP inhibitor verteporfin (VERT) at concentrations of 0.5, 1.0, and 2.0 µM for 24 h, in the absence of light, and the expression of nuclear SMAD 2/3 and YAP was resolved with fluorescent microscopy (**Figure** 4a).

**Figure 4.**
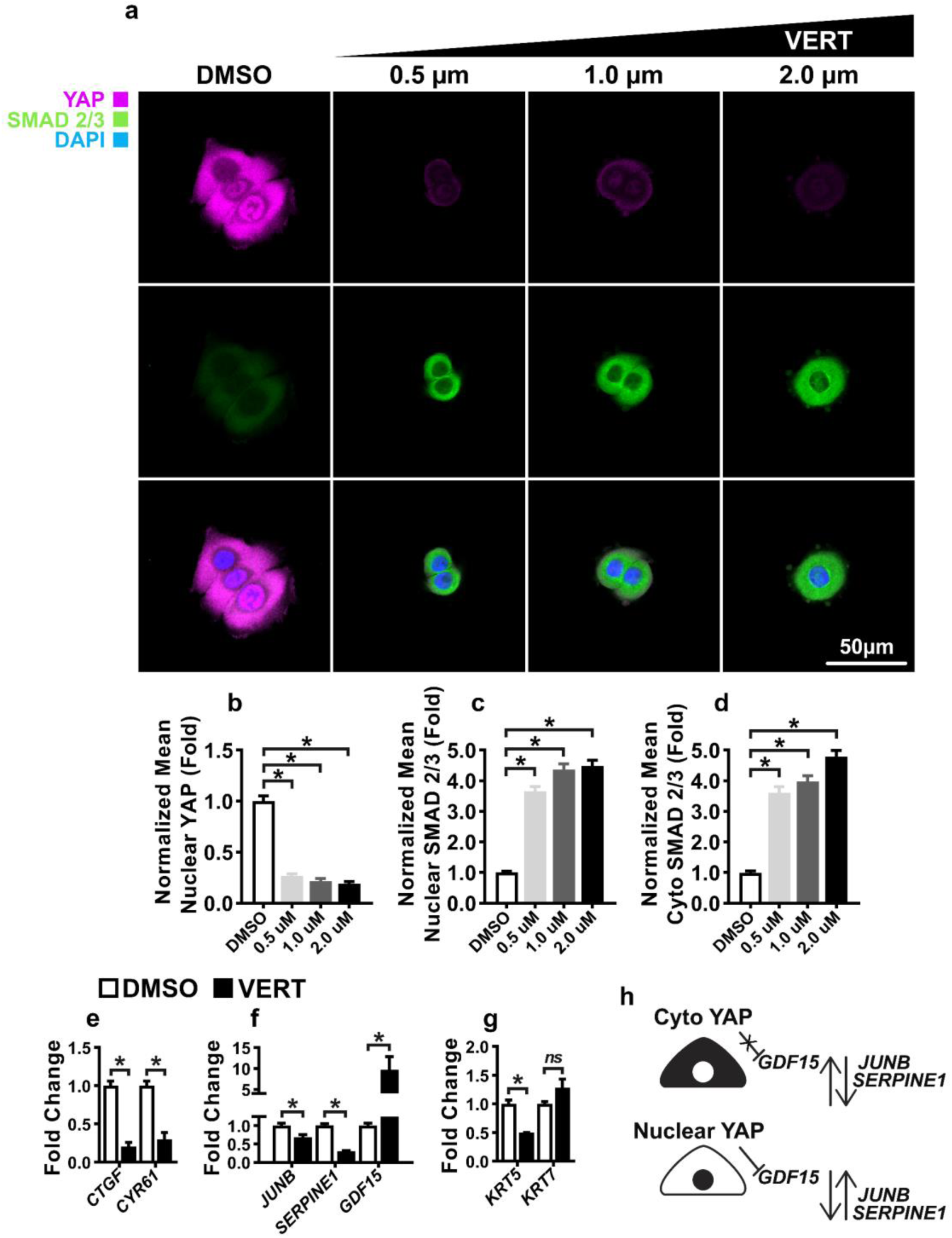
Loss of nuclear YAP stimulates *GDF15* expression and downregulates TGFβ target genes. **(a)** hS/PCs were cultured with verteporfin (VERT), 0.5, 1.0, and 2.0 µM, and a DMSO vehicle control for 24 h before SMAD 2/3 and YAP were visualized by ICC, nuclei were counterstained with DAPI. **(b-d)** Image quantification was performed with ImageJ to resolve nuclear YAP (b), nuclear SMAD 2/3 (c), and cytoplasmic SMAD 2/3 (d). Significance determined from one-way ANOVA followed by a Dunnett’s test, ^*^ indicates *p* < 0.05 relative to DMSO control. **(e-g)** hS/PCs were treated with 1 μM VERT for 24 h and mRNA express of YAP target genes: *CTGF, CYR61* (e); TGF-β targets genes: *JUNB, SERPINE1*, and *GDF15* (f); and keratins: *KRT5, KRT7* (g) were resolved with qPCR. (**h**) Schematic depiction detailing the role of nuclear YAP in supporting TGF-β signaling (*JUNB, SERPINE1*) and repressing *GDF15* expression. Error bars represent SEM in all cases. Two-tailed Student’s t-tests were conducted on data appearing in (e-g). ^*^ indicates *p* < 0.05 in all cases.

YAP expression decreased by ~0.75 fold with 0.5 μM of VERT, with further reductions in YAP observed in 1.0 and 2.0 µM VERT treated cultures (Figure 4b), indicating the proficiency of the VERT inhibition. Nuclear SMAD 2/3 was upregulated ~3.5 fold in response to 0.5 µM of VERT, and further enhanced as the concentration of VERT was increased (Figure 4c). However, the increased SMAD 2/3 observed with YAP inhibition was not concentrated to the cell nuclei, as cytoplasmic SMAD 2/3 levels were increased as well (Figure 4d). Importantly, these results demonstrate the inhibition of YAP in hS/PC cultures after VERT treatment.

To interrogate the effects of YAP inhibition on TGF-β signaling, TGF-β and YAP pathway transcripts in cultures receiving 1.0 µM of VERT were assessed with qPCR (Figure 4e-g). As expected, canonical YAP target genes *CTGF* and *CYR61* were downregulated, confirming the repression of YAP signaling with VERT treatment (Figure 4e).^[31]^ The TGF-β targets, *JUNB* and *SERPINE1*, were upregulated in hS/PC cultures after treatment with TGF-β1 (Figures S2b and S4a). YAP inhibition downregulated *JUNB* and *SERPINE1*, indicating the supporting role of YAP in the expression of these transcripts (Figure 4h). *GDF15* was modestly upregulated (1.5 fold, *p=0*.*08*) in TGF-β-1 treated cultures (Figure S3b); however, it was increased ~9 fold in response to VERT mediated YAP inhibition. Additionally, the role of YAP in the regulation of keratin expression was investigated (Figure 4g). *KRT5* was downregulated with the loss of YAP, but alterations in *KRT7* expression was not detected. These findings indicate that YAP represses *GDF15* expression while supporting the expression of the TGF-β signaling components *JUNB* and *SERPINE1* (Figure 4h).

### 2.5. TGFβR inhibition represses TGF-β1 induced K7 expression

After establishing that RGDSP cultures exhibited enhanced TGF-β1 expression and signaling, we investigated a strategy to mitigate TGF-β1 stimulated *KRT7*/K7 expression using the TGFβR inhibitor, A83-01. hS/PCs were cultured for 48 h with TGF-β1 and 2 µM of A83-01, a concentration reported not to disrupt BMP signaling,^[32]^ and characterized with fluorescent microscopy (**Figure** 5a).

**Figure 5.**
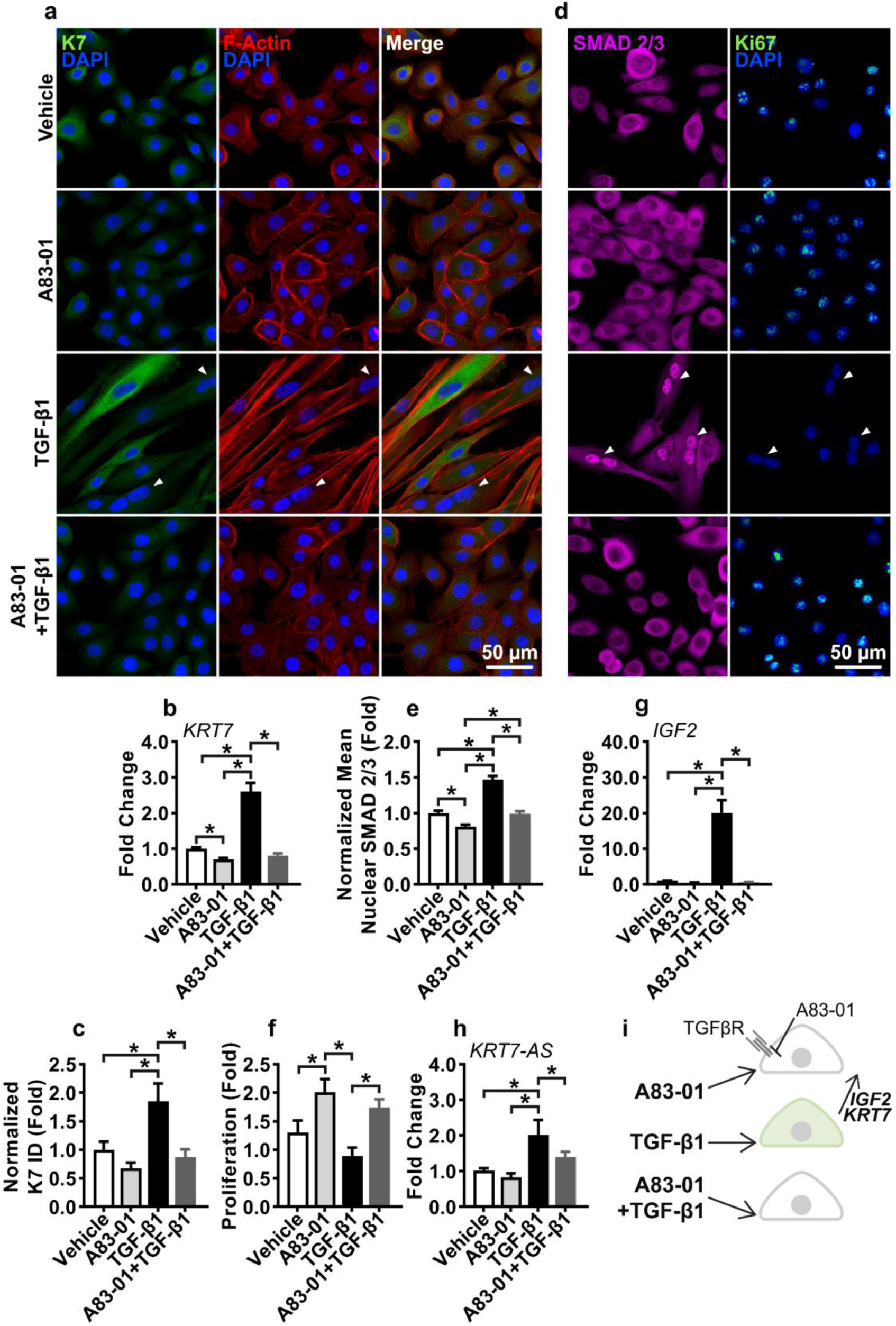
TGFβR inhibition represses TGF-β1 induced K7 expression. **(a)** hS/PCs were cultured with TGF-β1 and A83-01 for 48 h, and the expression of K7 and F-Actin was visualized with fluorescent microscopy. **(b-c)** A83-01 repressed TGF-β1 induced *KRT7* at the mRNA level (b) and at the protein level as determined by image analysis conducted with ImageJ (c). **(d)** hS/PCs were cultured with TGF-β1 and A83-01 for 48 h, and SMAD 2/3 and Ki-67 expression were investigated with ICC. White arrows indicate binuclear cells in TGF-β1 treated cultures. **(e)** ImageJ-derived analysis indicated TGF-β1 stimulated nuclear SMAD 2/3 that was inhibited by A83-01. **(f)** Proliferation was assessed by enumeration of DAPI stained nuclei in TGF-β1, and A83-01 treated cultures after 48 h of culture. **(g-h)** hS/PC expression of *IGF2* (g) and *KRT7-AS* (h) with TGF-β1 and A83-01 treatment were investigated with qPCR. Error bars represent SEM in all cases. One-way ANOVAs were performed on data presented in (b), (c), and (e-h) followed by Tukey’s multiple comparison test. ^*^ indicates *p* < 0.05 in all cases. **(i)** Schematic depiction of A83-01 inhibition of TGF-β1 stimulated K7 expression.

TGF-β1-treated cells were strikingly larger and extended with few cell-cell contacts, taking on a mesenchymal spindle shape morphology. A83-01 inhibition promoted an epithelial phenotype with F-actin localized to cell-cell contacts, and when provided in combination with TGF-β1, A83-01 prevented the TGF-β1-induced morphology changes. Immunofluorescence microscopy detailed a granular expression pattern of K7, which increased in response to TGF-β1 (Figure 5b). Furthermore, A83-01 was sufficient to resist the TGF-β1 mediated increase in K7 expression. Similarly, A83-01 inhibited the TGF-β1 stimulated *KRT7* expression while simultaneously decreasing *KRT7* expression below that of the vehicle control (Figure 5c). As TGF-β1 can utilize signaling pathways in addition to SMAD 2/3, which was elevated in RGDSP cultures, immunofluorescence microscopy was performed to demonstrate further that *KRT7*/K7 was correlated with the ability of TGF-β1 to induce nuclear SMAD 2/3 and inhibition of *KRT7*/K7 expression via A83-01 was also accompanied by reduced nuclear SMAD 2/3 (Figure 5d-e).

TGF-β1 is a cytostatic factor towards epithelial cells, and with TGFβR inhibition, provided by A83-01, hS/PCs responded with rapid proliferation (Figure 5f). In addition, competitively inhibited cultures receiving A83-01 retained higher proliferation than cultures receiving only TGF-β1. TGF-β1 did not reduce proliferation against the vehicle control over 48 h, but the non-G_0_ state cell cycle marker, Ki-67,^[33]^ was completely absent in TGF-β1 treated cultures (Figure 5d). However, nuclear Ki-67 was present in A83-01+TGF-β1 treated cultures, further demonstrating the ability of A83-01 to inhibit TGF-β signaling. Under the cytostatic effects of TGF-β1, cytokinesis failure can occur, producing binuclear cells,^[34]^ which were present in TGF-β1 treated cultures (Figure 5a, d). This is associated with genome instability, loss of imprinting (LOI), and led to the investigation of the expression of a long non-coding antisense (AS) RNA, *KRT7-AS*, reported to regulate *KRT7*/K7 expression.^[34-35]^ We determined the paternally imprinted gene of insulin-like growth factor 2 (IGF-2, *IGF2*) was ~20 fold upregulated in response to TGF-β1,^[27, 36]^ and concordantly, *KRT7-AS* expression was induced, (Figure 5g-h). Furthermore, TGF-β1 modestly upregulated insulin-like growth factor 1 receptor (IGF-1R, *IGF1R*), yet the expression of the non-imprinted insulin-like growth factor 1 (IGF-1, *IGF1*) was unaltered (Figure S6a-b).^[37]^ A83-01 medium supplementation prevented both TGF-β1 promoted *KRT7* and *IGF2* expression (Figure 5i), indicating that even SMAD-independent mechanisms would not escape inhibition by A83-01.

### 2.6. TGFβR inhibition represses RGDSP induced K7 expression

After associating nuclear SMAD 2/3 with increased *KRT7*/K7 expression and demonstrating that A83-01 can mitigate this response under 2D conditions, we investigated the effectiveness of A83-01 supplementation towards inhibiting TGF-β1 mediated K7 expression in RGDSP cultures. The sensitive TGF-β signaling balance maintained by hS/PCs was demonstrated when HA cultures without RGDSP were treated with A83-01 (**Figure** 6a). Proliferation, determined by DNA yield, in the HA+A83-01 cultures was reduced ~2 fold while a marginal increase in cell proliferation was observed in A83-01-treated RGDSP cultures (Figure 6b). While the proliferative properties of TGF-β inhibition were not maintained from 2D to 3D; *KRT7* and *FN* were repressed in A83-01 treated RGDSP cultures to the levels of the DMSO treated HA cultures (Figure 6c-d). Furthermore, we confirmed these findings by western blot and demonstrated that the RGDSP-promoted increase in K7 and fibronectin expression was repressible with inhibition of TGF-β signaling (Figure 6e). Although A83-01 treated HA cultures, with disrupted TGF-β signaling, exhibited increased *KRT7* and *FN* mRNA levels, increased protein levels of K7 and fibronectin were not detected (Figure 6e). On 2D substrates, we established the potential for epigenetic alterations to occur with TGF-β1 treatment of hS/PCs. When profiling the complete hydrogel culture time series, a distinct temporal event was found to occur at day 14, where *IGF2* was upregulated ~30 fold in 3D RGDSP cultures (Figure S6e). Yet, the expression of the *IGF1* or *IGF1R* genes did not follow (Figure S6f-g). As expected, inhibition with A83-01 prevented the *IGF2* increase in RGDSP cultures, and *IGF2* expression was not activated by inhibiting TGF-β signaling in HA cultures (Figure 6f). However, the enhanced expression of the *KRT7-AS* transcript that was observed in response to TGF-β1 was not induced in the RGDSP cultures, suggesting this mechanism is not required for the observed increased *KRT7*/K7 expression brought on by RGDSP (Figure 6g). qPCR and ICC indicated TGF-β inhibition allowed repression of K7 and maintenance amylase expression (Figure 6e-i). Furthermore, we found β-catenin remained localized to cell-cell junctions with TGF-β inhibition in RGDSP cultures (Figure 6i). Thus, we determined that the enhanced proliferation and amylase expression provided by RGDSP were maintained as K7 expression was suppressed with TGF-β signaling inhibition.

**Figure 6.**
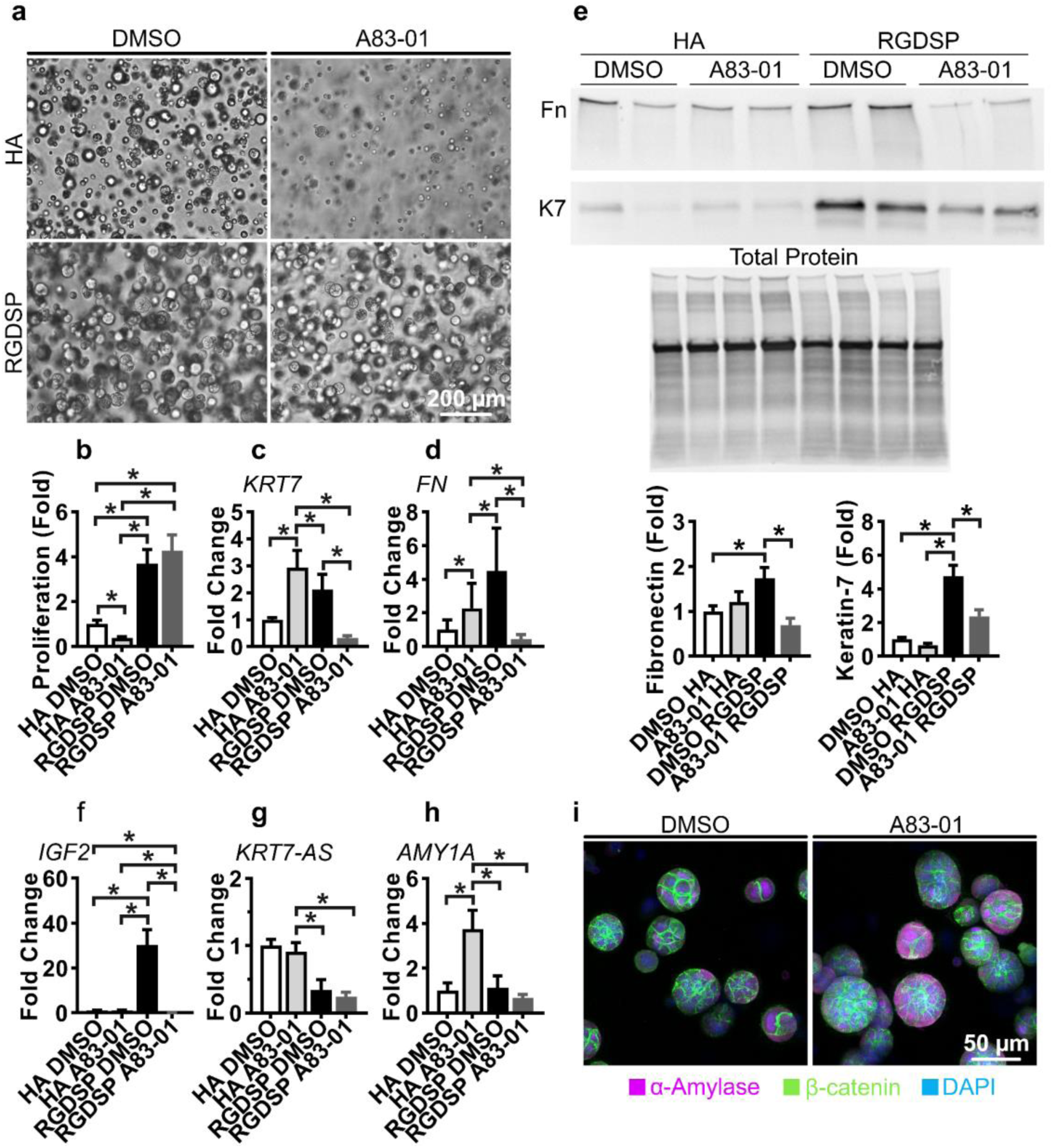
TGFβR inhibition represses RGDSP induced K7 expression. **(a)** hS/PCs were cultured in HA and RGDSP hydrogels with A83-01 or a DMSO vehicle control for 14 days and visualized with bright field microscopy. **(b)** dsDNA, indicative of proliferation, was resolved from simultaneous TRIzol™ RNA/protein extractions using the Quant-iT™ PicoGreen™ assay. **(c-h)** hS/PCs were cultured in HA and RGDSP hydrogels for 14 days receiving A83-01 or DMSO, and expression of *KRT7* (c), *FN* (d), *IGF2* (f), *KRT7-AS* (g), and *AMY1A* (h) were assessed with qPCR. **(e)** hS/PCs were cultured in HA and RGDSP hydrogels for 14 days receiving A83-01 or DMSO, and western blotting was conducted to investigate K7 and fibronectin expression. Duplicate measurements are presented from technical replicates extracted from separate hydrogels. **(i)** hS/PCs were cultured for 14 days in RGDSP receiving A83-01 or DMSO, and fluorescent microscopy indicated that expression α-amylase and β-catenin was not suppressed by treatment with A83-01. Error bars represent SEM in all cases. One-way ANOVAs were performed on data presented in (b-h) followed by Tukey’s multiple comparison test. ^*^ indicates *p* < 0.05 in all cases.

## 3. Discussion

We report here that an RGDSP peptide stimulates K7 expression in human salivary gland cells. We discovered that K7 expression was accompanied by the expression of TGF-β superfamily members GDF-15 and TGF-β1. TGF-β1, but not GDF-15, was capable of initiating K7 expression and SMAD 2/3 signaling. Loss of nuclear YAP was found to increase GDF-15 expression and impair TGF-β signaling. However, the loss of nuclear YAP was insufficient to shut down TGF-β signaling, and nuclear SMAD 2/3 was amplified without the aid of nuclear YAP in RGDSP cultures. TGFβR inhibition suppressed K7 expression initiated from both RGDSP hydrogel culture and exogenously supplied TGF-β1 in 2D environments. Though ductal cells are characterized by K7 expression, these findings suggest that K7 expression can be stimulated under conditions that induce TGF-β1.

This work assessed the temporal dynamics of transcription, protein expression, signaling, and morphological characteristics of hS/PCs cultured in RGDSP-hydrogels, providing insight into the resulting K7-expressing phenotype and the factors behind its emergence. We demonstrate the ability of TGF-β1 to regulate K7 expression in 2D and 3D environments, indicating its dominant role behind the expression of K7, irrespective of a cell’s current environment, morphology, cell cycle, or cytokine profile. We describe the initiation of K7 from a non-terminal differentiation point maintained by progenitor cultures, yet the activation of K7 expression is similar to the events described to occur in differentiated acinar cells.^[1a, 6a, 6c, 6d, 6f]^ As we did not perform this work from the onset of cell isolation from human tissues, we were unable to demonstrate recovery of acinar markers, such as Mist1/*BHLHA15* expression, with TGFβR inhibition. Furthermore, these findings suggest that TGFβR inhibition may in some cases be deleterious and that a degree of SMAD 2/3 signaling is required to sustain SG cell cultures.

In RGDSP cultures, nuclear SMAD 2/3 increased with culture time, while nuclear YAP decreased (Figure 3c-e). We found that YAP localization played a central role in modulating both TGF-β signaling and GDF-15 expression. The ability of YAP to potentiate SMAD signaling when located in the nucleus and act as an inhibitor of SMAD signaling when located in the cytoplasm has been reported in a multitude of biological models.^[19a, 20, 38]^ In agreement with those findings, we found pharmacological YAP inhibition decreased TGF-β target genes (Figure 4f), demonstrating that nuclear YAP can support TGF-β signaling. However, we found that the loss of nuclear YAP was not sufficient to suppress nuclear SMAD 2/3 accumulation. This could be due to the fact that the loss of nuclear YAP also resulted in the downregulation of cytoplasmic YAP expression (Figure S4a), thus decreasing the pool of cytoplasmic YAP available to suppress SMAD signaling and allowing aberrant SMAD signaling to proceed. Additionally, the regulation of GDF-15 expression by YAP is supported by prior work, where genetic overexpression, mechanical inactivation, and VERT inhibition have demonstrated the ability of YAP to suppress GDF-15.^[39]^ While we cannot correlate *GDF15* mRNA expression with nuclear YAP levels in RGDSP cultures (Figure S4c), protein level GDF-15 expression was well correlated (Figure 3a, c). The mechanisms underlying GDF-15 expression are not well understood, increased protein levels have been reported in the absence of increased mRNA expression, and post-transcriptional control mechanisms have been described.^[40]^ Thus, we predict YAP maintains control of GDF-15 expression in RGDSP cultures despite the observed mRNA expression profile of *GDF15*.

GDF-15 is in some cases reported to use SMAD signaling pathways and could be theorized to produce the increased SMAD 2/3 we observed with YAP inhibition.^[40-41]^ However, we were unable to visualize increased SMAD 2/3 levels in hS/PCs exposed to recombinant GDF-15 (Figure S3d). However, exogenous GDF-15 induced *IL1B* expression (Figure S3b), suggesting the supplied GDF-15 retained biological activity. VERT was initially developed for use in photodynamic therapy, yet its non-photoactivated actions have made it the most well recognized molecular inhibitor of YAP-TEAD transcription. Despite this, in the absence of light, VERT inhibits not only YAP but also autophagy.^[42]^ SMAD 2/3 levels are maintained by proteasomal degradation;^[43]^ however, disrupted autophagy leads to protein accumulation and a decreased rate of total protein turnover.^[43-44]^ Additionally, F-actin maintains autophagic flux,^[42a]^ and we observed that VERT strongly disrupted the F-actin of hS/PCs (Figure S5). Thus, we anticipate that the additional autophagy inhibition properties of VERT contributed to the increased SMAD 2/3 levels observed in our cultures.

It might appear contradictory that the adhesive RGDSP peptide downregulates the expression of YAP, but this is predictable due to the non-degradable nature of the hydrogel. A recently proposed model of YAP regulation in 3D matrices indicates that ‘conforming properties’ of a hydrogel network are required to sustain traction forces for YAP signaling; without stress relaxation or pericellular degradability, progressive compressive confinement leads to contact inhibition as cell division proceeds.^[19a]^ In addition to the regulation of YAP localization by substrate stiffness, 2D studies report that YAP localization is dictated by the isoform-specific integrin affinity and ability to provide integrin-mediated traction forces from an ECM substrate.^[24b, 45]^ The major mammalian hyaluronidases are Hyal-2 and Hyal-1; Hyal-2 can degrade high molecular weight HA to 20 kDa fractions, and Hyal-1 can reduce HA to tetrasaccharides; however, it requires acidic conditions and is limited to a minimum HA substrate of hexasaccharide size.^[46]^ After these synthetic HA hydrogels are formed, disulfide crosslinking continues, quickly placing a covalent crosslink on each hexasaccharide HA unit, and after ~83% of the thiols are consumed, approximately 7 days, a crosslink exists on every other disaccharide.^[21]^ Due to the extent of crosslinking and limited ester hydrolysis exhibited by these hydrogels, (~2% by mass), in this context, these hydrogel networks can be considered non-degradable.^[21]^ Therefore, as the encapsulated hS/PCs proliferate to form multicellular structures, they are required to push away the HA matrix to accommodate their increasing volume. As a result of this displacement, the HA matrix reciprocally increases a compressive, YAP deactivating, pressure against the cells.^[47]^ Furthermore, we have recently demonstrated that RGDSP peptides promote hS/PC attachment and spreading on these hydrogel matrices that were repressed by treatment with a β1-integrin blocking antibody.^[23]^ However, cells encapsulated in control HA hydrogels are only capable of binding HA through their CD44 or RHAMM receptors and would require *de novo* synthesis of ECM to initiate integrin activation.

Incorporating integrin affinity into our 3D model YAP signaling, in the single-cell state at day 1, hS/PCs bind the RGDSP peptide, activating mechanoresponsive YAP signaling, which correlates with *YAP1, CTGF*, GDF-15 expression, and filopodia extension. We did not investigate if the filopodia mechanically induced nuclear YAP localization or a product of YAP signaling. However, inhibition with VERT strongly disrupted F-actin polymerization, suggesting that YAP maintains actin polymerization. Additionally, F-actin is a known target of YAP signaling,^[19a, 20]^ and YAP deletion in murine submandibular glands disrupted F-actin expression and organization.^[48]^ As cell division led to increased cell-cell adhesion, depicted by the β-catenin localization in (Figure 1f), contact inhibition and compressive forces supplied by the hydrogel network mechanically inactivated YAP and stimulated the expression of GDF-15. A similar temporal profile of YAP localization has been experimentally reported for iPSC organoids cultured in stiff degradable matrices as described by the ‘conforming properties’ model of 3D YAP localization.^[19a, 49]^

Basal expression of GDF-15 is low in most tissues, and it is only expressed during states of cellular stress.^[11b, 11c, 40]^ In RGDSP cultures, nuclear YAP was downregulated with spheroid development, and this presented as SASP with high GDF-15 and MMP-1 expression (Figure 3a, Figure S3c). We investigated a requirement for senescence or SASP with K7 expression, yet RGDSP cultures did not maintain expression of the senescence hallmarks *CDKN1A* and *CDK2A* (Figure S3a, b). Thus, we could not confirm a role for senescence or SASP towards developing the K7 expressing phenotype, but the loss of nuclear YAP explains the elevated GDF-15 expression.

Our work agrees with previous reports, showing that stress induces keratin expression of primary salivary gland cells and loss of the acinar phenotype.^[6a]^ Studies performed with freshly isolated murine SG cells have reported the increased expression of keratins to include K5, K7, and K19. Here, we observed differential regulation of these keratins. Human SG progenitor cells express high levels of K5 at passage 3,^[50]^ and we found K5 expression was maintained but not upregulated while K7 expression increased in RGDSP cultures (Figure 2b). *KRT18* and *KRT19* were also not upregulated with *KRT7*/K7 expression (Figure S2c), suggesting a separate keratin may be dimerizing with K7 in these cultures. More so, 2D cultures treated with TGF-β1 moderately decreased, (0.53 fold; *p=0*.*077*), *KRT5* mRNA (Figure S6d). Our findings are supported by literature detailing the independent regulation of keratin subtypes.^[28]^ Additionally, we also describe K7 expression after an initial stage of increased K5 expression occurring in isolation and passage.^[50]^ Further work is required to determine if keratin subtypes are independently regulated during the increased keratin expression occurring at isolation.

Amylase expression was increased in RGDSP cultures expressing K7 (Figure 2a-c). *In-vivo*, amylase expression is unique to acinar cells and not expressed by ductal cells.^[27]^ Although amylase is expressed by acinar cells, Mist1 expression is required to specify acinar identity.^[1a]^ Mist1 levels remain low in our cultures after isolation,^[50]^ and transcription of NKCC1 (*SLC12A2*), typically expressed by acinar cells, continuously decreases in the RGDSP cultures (Figure 2a). In agreement with our findings, Mist1 and NKCC1 expression is reported to decrease rapidly during *in-vitro* SG culture, but the impact on amylase expression is less severe.^[1a, 50-51]^ Here, we report that this RGDSP culture system can increase amylase expression; this is substantiated by previous reports of ductal derived stem/progenitors cells directed to increase amylase expression by culture-imposed differentiation.^[2, 52]^

Although TGF-β1 is pro-fibrotic and detrimental to SG function,^[14, 19b, 53]^ studies performed with *in*-*vitro* cultured SG cells, have not reported conclusive evidence that exogenous TGF-β1 is deleterious to amylase expression or secretion.^[8a, 54]^ The ability of hS/PCs to form organized multicellular structures with adherins junctions is expected to contribute to the expression of amylase seen in RGDSP cultures. These findings suggest, separate regulation of amylase from acinar or ductal lineages, and further work is required to characterize this phenotype. Importantly, amylase expression was maintained with TGFβR inhibition indicating amylase expression could be separated from the K7 phenotype, therefore increasing the acinar characteristic of the RGDSP cultures.

Here, we found that TGF-β1 can stimulate K7 expression and that the TGFβR inhibitor, A83-01, can inhibit TGF-β1 dependent K7 expression (Figure 5B). Media supplementation with TGF-β/BMP/SMAD inhibitors has found broad applicability in primary epithelial cell culture for their ability to prevent epithelial differentiation and senescence.^[8a, 8b, 55]^ In the SG, TGFβR inhibition increased acinar marker expression in studies using passage 9 primary murine submandibular gland cells.^[8a]^ When murine submandibular cells were isolated and maintained with a TGFβR inhibitor, an enriched progenitor phenotype that exhibited decreased K7 and aquaporin-5 (acinar marker) expression was observed.^[8b]^ However, a TGF-β1 dependant role for the induction of K7 expression in the SG has yet to be confirmed. In ovarian cancer cells, a similar TGF-β1-SMAD 2/3-K7 mechanism has been described.^[9e]^ More broadly, TGF-β superfamily members regulate K7 expression across various tissues.^[9]^ Thus, our findings are novel and supported by the known role of TGF-β signaling in the regulation of epithelial differentiation and K7 expression.

Our findings suggest that the stress imposed by TGF-β1 can induce features of EMT and epigenetic aberrations in SG cells alongside K7 expression. In the SG, chronic exposure to TGF-β is repeatedly reported as deleterious to SG function; however, attention has primarily been directed to the fibrotic tissue that accumulates and displaces glandular epithelium.^[14, 19b, 53]^ We report that TGF-β1 treated hS/PCs in 2D cultures take on a morphology indicative of a mesenchymal phenotype (Figure 5a, d). This is supported by literature reporting increased expression of the mesenchymal marker vimentin with K7 during the isolation and culture of primary SG cells.^[6c, 6d]^ During EMT, epigenome maintenance can become compromised, activating otherwise repressed genes.^[17]^ TGF-β1 can initiate epigenetic events by altering chromatin remodeling, histone acetylation, DNA methylation, and activating the expression of long non-coding RNAs (lncRNA).^[56]^ We found that TGF-β1-stimulated *KRT7-AS* and *IGF2* expression (Figure 5g, h), which were repressed by A83-01 in 2D cultures. While *IGF2* performed similarly in RGDSP cultures, we could not confirm a role for *KRT7-AS* in 3D cultures (Figure 6f, g). However, the unique expression of *IGF2* with *KRT7* in RGDSP cultures without *IGF1*, or *IGF1R* expression further suggests the occurrence of an epigenetic event. *IGF2* is a parentally imprinted gene whose expression is tightly regulated and only weakly expressed in the healthy adult SG.^[27, 36]^ However, *IGF2* is highly expressed in various cancers due to epigenetic activation of the repressed maternal allele.^[57]^ We did not investigate the mechanism of *IGF2* expression; however, in multiple reports, TGF-β1 has been shown to activate *IGF2* expression, as well as the co-imprinted gene, *H19*.^[58]^ To our knowledge, exogenous TGF-β1 has not been demonstrated to alter imprinting status. Nevertheless, imprinting status does not correlate with *IGF2* expression in iPSC cultures, and chromatin accessibility is the dominant regulator of *IGF2* expression.^[59]^ Interestingly, a SMAD 3-CCCTC-binding factor (CTCF) complex has been proposed to regulate expression at the *IGF2* locus by restructuring chromatin to increase crosstalk,^[58d]^ providing a plausible mechanism for TGF-β regulation of IGF-2. The expression of *KRT7-AS* and *IGF2* oncogenes with *KRT7*/K7 further indicates that the K7 phenotype is a response to cytokine-induced stress. Though EMT-associated epigenetic events were found with K7 expression in RGDSP cultures, cell-cell contacts were maintained, β-catenin was localized to adherins junctions, and filopodia regressed (Figure 1d), which are each characteristic of an enhanced epithelial phenotype.

We have reported that the ductal marker K7 co-expressed with features observed during EMT in response to TGF-β1. While we do observe similarities to the ductal keratin expression observed in injured de/transdifferentiating pancreatic acinar cells,^[60]^ overexpression of intermediate filaments (keratin, desmin, nestin, and vimentin), is a broadly recognized stress response.^[61]^ Intermediate filaments not only impart mechanical support after injury, but they can also provide anti-apoptotic signals by acting as a sponge for aberrant phosphorylation, providing a scaffold for 14-3-3 proteins, as well as sequestering apoptotic proteins.^[61a, 62]^ Thus, there is strong evidence that K7 expression indicates a state of cellular stress; however, we cannot conclude this indicates ductal de/transdifferentiating, and further research is required to determine its relationship with EMT. Our findings suggest TGF-β1 is involved in the loss of acinar markers during primary SG cell culture and is supported by prior literature.^[6b-d, 8b]^ We conclude that RGDSP stimulated TGF-β1 or exogenously supplied TGF-β1 can induce K7 expression of hS/PCs.

TGF-β1 levels slowly rise in HA cultures over time, suggesting TGF-β1 expression is inherent to spheroid development, with or without the supplied integrin adhesion. Nevertheless, literature suggests the inflammatory properties of RGDSP can be mitigated by including its synergy sequence to target α_5_ as opposed to α_V_ integrin subunits.^[63]^ Also, incorporating dual active and passive degradation mechanisms has successfully mitigated the inflammatory response arising from synthetic hydrogel culture.^[49]^ Although imparting enzymatic degradation in synthetic hydrogel designs has been beneficial towards sustaining an acinar phenotype, it is only partially sufficient, and the underlying problem has not been identified.^[6e, 51b]^ The hydrogel platforms used here are subject to few temporal factors allowing us to precisely deconstruct the signaling behind K7 expression. Our work will be realized in future hydrogel designs by incorporating properties to negate the signaling and overexpression of TGF-β1.

It remains to be seen if TGFβR inhibition applied at the point of cell isolation can prevent K7 expression and the loss of Mist1 in primary human SG cultures. TGF-β1 suppresses inflammatory responses and BMP signaling, and our work suggests that balanced SMAD signaling is required to sustain SG cultures.^[18, 19b]^ We expect incorporating other known proacinar factors, FGF2, FGF7, FGF10, laminin-111, and Y-27632, will be further beneficial towards suppressing ductal differentiation and maintenance of the acinar phenotype.^[6e, 22b, 22d]^

## 4. Conclusion

In human salivary gland progenitor cells, TGF-β1 can induce K7 expression in a SMAD2/3 dependent manner which is suppressible by a TGFβR inhibitor. When hS/PCs are encapsulated in RGDSP presenting hydrogels, SMAD 2/3 signaling proceeds independently of nuclear YAP expression and is maintained by TGF-β1 levels. Expression of K7 by adult SG cells is a TGF-β1 dependent stress response that can involve epigenetic aberrations with features of EMT.

## 5. Experimental Section

### Materials

Reagents were procured from Fisher Scientific and used as received unless otherwise noted.

### Cell Isolation and Maintenance

hS/PCs were isolated from human salivary gland tissue, and cultured following reported procedures.^[21, 50]^ Tissue was obtained from consenting patients in agreement with protocols approved by institutional review boards at Christiana Care Health Systems and the University of Delaware. hS/PCs were maintained in HepatoSTIM™ medium (355056; Corning Inc., Corning, NY) supplemented with 100 U mL^−1^ penicillin-streptomycin, 1% (v/v) amphotericin B, and 10 ng mL^−1^ epidermal growth factor (EGF) (Corning Inc.). Passaging was conducted at 70-80% confluence using 0.05% (w/v) trypsin-EDTA. Trypsin was neutralized using a trypsin soybean inhibitor (T6522; Sigma Aldrich, St. Louis, MO). Experiments were conducted with at least three different donors at passages between 3-4.

### Hydrogel Synthesis

Thiolated and acrylated HA derivatives (HA-SH and HA-AES) and maleimide-functionalized RGDSP (MI-RGDSP) were synthesized as previously reported.^[21, 64]^

### 3D hS/PC Encapsulation in HA Hydrogels

HA-SH, 1% (w/v), was reconstituted in sterile PBS to prepare HA hydrogels. To prepare RGDSP hydrogels, a 0.25 mM solution of MI-RGDSP in PBS was sterilized by filtration through a 0.2 μm Acrodisc® PTFE syringe filter (4602; Pall Corporation, Port Washington, NY). The resulting HA-SH or HA-SH/RGDSP solutions were neutralized to pH 7.4 using sterile 0.1 M NaOH, and hS/PC cell pellets, targeting a final cell concentration of 1 × 10^6^ cells mL^−1^ were suspended in these solutions by gentle pipetting. The HA-SH laden cell suspensions were combined with HA-AES at a 1/20 (v/v) ratio to initiate crosslinking. Characterization of protein and mRNA expression was conducted using 100 μL hydrogels prepared in 12 mm Millicell® 04 μm PTFE cell culture inserts (PICM01250; EMD Millipore) receiving 800 μL HeptoSTIM™ media every 72 h. ICC characterization was conducted using 30 μL hydrogels cultured in 48-well uncoated glass-bottom plates, 1.5 coverslip, (P48G-1.5-6-F; MatTek, Ashland, MA) receiving 300 μL of fresh HeptoSTIM™ media every 72 h. When A83-01 inhibition was performed, 2 uM A83-01 in HeptoSTIM™ and DMSO supplemented media were exchanged every 48 h.

### 2D hS/PC Cell Culture and Characterization

hS/PCs were seeded at 35,000 cells cm^−3^ and cultured for 48 h before introducing the indicated growth factors and inhibitors. For ICC studies, cells were cultured on 8-well Nunc™ Lab-Tek™ II chambered coverglass (155409; Thermo Scientific™, Waltham, MA). For gene expression studies, cells were cultured in Nunc™ cell-culture treated 6-well plates (140675; Thermo Scientific™). HepatoSTIM™ medium was supplemented with 10 ng mL^−1^ of recombinant human TGF-β1 (100-21; PeproTech®, Rocky Hill, NJ), 100 ng mL^−1^ of recombinant human GDF-15 (120-28C; PeproTech®), and 2 µM of A83-01 (9001799; Cayman Chemical Company, Ann Arbor, MI) as single factors or in combination when appropriate. Verteporfin (VERT) (SML0534; MilliporeSigma) ICC studies were conducted at 0.5, 1.0, and 2.0 µM, in complete darkness,^[65]^ and cultures were terminated after 24 h of culture. Gene expression studies were performed with VERT at 1.0 µM. In all cases, vehicle controls were maintained using PBS, DMSO, or a combination thereof, as required. RNA isolation was conducted with TRIzol™ and purified with RNA Clean & Concentrator™-5 Kit as described in the 3D isolation protocol.

### 3D Immunofluorescence

Hydrogel constructs were fixed with 4% (w/v) paraformaldehyde (PFA) in PBS for 1 h. PBS was supplemented with 500 U mL^−1^ penicillin-streptomycin and 0.05% (v/v) sodium azide to produce PBS+PS, which was combined with the permeabilization agent detailed in Table S2 to prepare PBS/perm. Blocking was conducted for 16 h at 4 °C with 3% (w/v) bovine serum albumin (BSA) reconstituted in PBS/perm (BSA/perm). Primary antibodies were diluted in BSA/perm, and a 48 h incubation was performed on the fixed constructs. Hydrogel constructs were then washed with PBS/perm that was exchanged 5 times over 24 h. Alexa Fluor® AffiniPure Fab Fragment Anti IgG (H+L) secondary antibodies (Jackson Immunoresearch Labs, West Grove, PA) were diluted at 1/200 (v/v) in the BSA/perm solution, and constructs were subsequently incubated in the secondary antibody solution for 48 h at room temperature. Hydrogel constructs were then washed with PBS/perm that was exchanged 3 times over 3 h. Next, 4′,6-diamidino-2-phenylindole (DAPI) (D1306; Life Technologies, Carlsbad, CA) and Alexa Fluor™ 568 Phalloidin (phalloidin) (A12380; Life Technologies, Carlsbad, CA) were diluted at 1/1000 and 1/450 (v/v) respectively in PBS/perm. Constructs were incubated in DAPI and phalloidin solutions for 12 h at room temperature. The hydrogel constructs were then washed for 6 h with PBS/perm that was exchanged 3 times. ICC incubation and wash steps were conducted using an orbital shaker at 250 RPM. Mounting was conducted by incubating for 16 h at 4 °C with VECTASHIELD® PLUS Antifade Mounting Medium (H-1900; Vector Laboratories, Burlingame, CA). Extended information detailing primary antibody, secondary antibody, and permeabilization agent can be found in Table S2. Fluorescent microscopy was performed with a Zeiss LSM 880, ZEN 2.3 SP1, equipped with an Airy scan super-resolution detector (Carl Zeiss, Oberkochen, Germany). An LD LCI Plan-Apochromat 25×/0.8 Imm Korr objective was used with Immersol™ 518F (Carl Zeiss), refractive index 1.518. Image acquisition was performed using Fast Airyscan mode, and a piezoelectric stage was used to sequentially capture single-channel excitations along the z-axis. Volumetric images were recorded with an x-y area of 83.54 µm and 0.492 µm z-axis scaling. Fast Airyscan processing was performed with ZEN 3.0 SR software producing 16-bit images, and SMAD 2/3 and YAP expression was quantified using Imaris 9.7.0 3D-4D Imaging Software (Oxford Instruments, Abingdon, United Kingdom). Maximum intensity projections of the F-actin channel were prepared using ImageJ software (NIH, Bethesda, MD, https://imagej.nih.gov/) and the ImageJ plugin FiloQuant (https://imagej.net/plugins/filoquant) was utilized to analyze filopodia length.^[66]^ Similarly, maximum intensity projections of the β-catenin channel was generated with ImageJ, and the radial distribution of β-catenin was assessed using the radial profile plugin. (https://imagej.nih.gov/ij/plugins/radial-profile.html) Samples were normalized relative to their maximum radial profile intensity before measurements were averaged.

Following scientific digital image ethical guidelines,^[67]^ representative hS/PC structures were centered to the field of view by cropping the x-y area to 63.39 µm before uniformly applying brightness adjustments with ZEN 3.0 SR software. To minimize power variation and laser instability during quantitative imaging, imaging was completed within 3 sessions, and lasers were warmed for a minimum of 30 min prior to the start of an image session.^[67a]^ Pilot studies indicated that quantitative fluorescent intensity measurements were valid at distances, z-dimension, up to 500 μm into the hydrogel constructs, and image collection was limited to this distance. Briefly, Alexa Fluor® 647 labeled reference standards (BLI887A-1; Polysciences, Warrington, PA) were encapsulated in HA hydrogels, and z-stack images were collected from the coverslip to the working distance of the piezoelectric stage. Imaris Imaging Software was used to derive the reference standard mean intensities, and their corresponding distance from the coverslip was fit to a linear model (Figure S1).

### 2D Immunofluorescence

hS/PC cultures were terminated with PFA for 30 min. PBS+PS was used to reconstitute BSA with the appropriate permeabilization agent as detailed in Table S4 to prepare PBS/perm. Blocking was conducted for 16 h at 4 °C with the indicated (Table S4) PBS/perm solutions. Primary and highly cross-adsorbed whole goat IgG Alexa Fluor® (A-11029, A-21236, A-11034, A-21245; Invitrogen™) secondary antibodies were incubated, sequentially, for 16 h at 4 °C followed by 1 h at room temperature. Secondary antibodies were diluted at 1/250 in all cases. Following antibody incubations, samples were washed with PBS/perm that was exchanged 6 times over 6 h at room temperature. Next, DAPI and phalloidin were diluted at 1/1000 and 1/450 (v/v) respectively in PBS/perm and incubated with samples for 90 min at room temperature. Samples were mounted with VECTASHIELD® PLUS antifade mounting medium. Extended information detailing primary antibody, secondary antibody, and permeabilization agent can be found in Table S4.

Fluorescent microscopy was conducted with a Zeiss LSM 880, ZEN 2.3 SP1, fitted with an Airy scan super-resolution detector. Imaging conducted with C-Apochromat 10×/0.45W or LD LCI Plan-Apochromat 25×/0.8 Imm Korr objectives were used with deionized water or Immersol™ 518F (Carl Zeiss) respectively. Image acquisition was conducted using Fast Airyscan mode and with sequential channel capture, frame mode. Fast Airyscan processing was performed with ZEN 3.0 SR software, generating 16-bit images, and quantification of protein expression and localization was assessed with ImageJ. Binary nuclear images were produced by applying a moments threshold to the DAPI channel, then a watershed segmentation and adding to the ImageJ ROI manager using the magic wand tool. The corresponding cell bodies were isolated by manually tracing the F-actin channel using the segmented line tool and adding the selection to the ROI manager. A Huang threshold was applied to the SMAD2/3 channel, and individual cell measurements were made using the limit to threshold function. Cytoplasmic expression was determined by subtracting the integrated density (ID) of nuclear measurements from the corresponding cell body ID measurements. ID is reported after background corrections were applied using the relationship (ID = area × mean gray value) and normalizing to the mean ID of non-treated controls from each biological experiment. K7 expression was analyzed similarly as above; segmentation was performed by manually tracing the F-actin channel using the segmented line tool, and a Huang threshold was applied to the K7 channel. Brightness and contrast adjustments were uniformly applied to the selected representative images using ZEN 3.0 SR software. A blinded reviewer performed analysis, and a minimum of 60 measurements was made for each condition.

### Immunoblot Array

Expression of TGF-β family members was investigated using a Human TGF beta Array C2 (AAH-TGFB-2-2; RayBiotech®, Peachtree Corners, GA). Medium collected from hydrogel cell constructs from days 3-6 was pooled from three biological replicates and used to carry out the assay following the manufacturer’s procedure. Chemiluminescent signals were recorded using an iBright™ FL1500 Imaging System.

### Enzyme Linked Immunosorbance Assay (ELISA)

Medium was collected from hydrogel cell constructs on days 3, 6, 9, and 12 of culture and stored at −80 °C until analysis. Total TGF-β1 levels were assessed using Human TGF-beta 1 DuoSet® ELISA (DY240; R&D Systems™, Minneapolis, MN). Similarly, GDF-15 expression was analyzed using Human GDF-15 DuoSet® ELISA (DY957; R&D Systems™). For the activation of latent TGF-β1, the medium was treated with Sample Activation Kit (DY010; R&D Systems™). TGF-β1 and GDF-15 ELISAs were performed using Substrate Reagent Pack (DY999; R&D Systems™), and absorbance measurements were made at 450 nm with a 570 nm correction using a SpectraMax® i3x Multi-Mode Microplate Reader. α-Amylase expression was quantified using the Human Salivary Amylase Alpha ELISA Kit (NBP2-68203; Novus Biologicals™, Littleton, CO), and luminesce measurements were made using a SpectraMax® i3x Multi-Mode Microplate Reader. All assays were performed in accordance with the manufacturers’ protocols. Protein levels were normalized to the number of cells in each hydrogel at each time point using the procedure detailed in the 3D Proliferation method conducted on days 1, 3, 6, 9, and 12 of culture.

### 3D Cell Proliferation

hS/PC proliferation in 3D constructs was determined by enumerating nuclei in confocal z-stack images collected on days 1, 3, 7, and 14 of culture. Cell-laden hydrogel constructs were fixed with 4% PFA at the designated time points, and nuclei were labeled with Hoeschst 3342 (hoeschst) (ThermoFisher Scientific) at 10 μg mL^−1^ in PBS for 30 min. Microscopy was conducted with a Zeiss LSM 880 equipped with an Airyscan detector using a 10×C-Apochromat 0.45 N.A water immersion objective. Images were captured as 300.45 μm z-stacks with a 1.757 μm z-axis step size and x-y area of 850.19 µm, and fast airy scan processing was performed using Zen Black 3.0 SR. Nuclei were resolved and counted using 3D-4D Imaging Software Imaris 9.7.0. Three independent experiments were conducted at each time point.

### Simultaneous RNA, DNA, and Protein Isolation from 3D Cultures

Isolation of protein and nucleic acids from the same experimental sample was conducted using TRIzol™ (15596026; Invitrogen, Carlsbad, CA) and adapting the manufacture’s recommended procedure. Detailed methodology of nucleic acid and protein extraction is included in the supporting information. Additional information regarding RT-qPCR, Quant-iT™ PicoGreen™ dsDNA Assay Kit (P11495; Invitrogen™), and western blot assay conducted with chemifluorescencent detection (ECF™) are included in the supporting information.

### Statistical Analysis

Comparisons between two experimental groups were made using a two-tailed Student’s t-test. A one-way analysis of variance (ANOVA) was performed when three or more experimental groups were compared, and a two-way ANOVA was performed when data sets included an additional variable. When ANOVAs returned an F statistic greater than F critical, a post-hoc Tukey multiple comparisons test was carried out. For multiple comparisons containing a control group, a post-hoc Dunnett’s test was performed. *p* < 0.05 was deemed significant and indicated by ^*^,^†^,^‡^, or ^§^. Significance from day 1, 3, 6 and 7, 9 and 12 samples of the same experimental group are indicated by ^†^,^‡^, and ^§^ respectively. Statistical interpretations were made using JMP Pro 15 (SAS Institute Inc., Cary, NC).

## Acknowledgments

This work was supported in part by the National Institutes of Health (NIDCD, R01DC014461; NIDCR R01 DE029655), National Science Foundation (NSF, DMR 1809612), and Delaware Bioscience Center for Advanced Technology (DE-CAT 12A00448). EWF acknowledges the University of Delaware for the Dissertation Fellowship. Instrumentation support was made possible by NIH (S10 OD016361, P20 GM103446) and NSF (CHE-0840401, CHE-1229234, IIA-1301765, DMR-2011824) grants. Microscopy access was supported by grants from the NIH-NIGMS (P20 GM103446), the NSF (IIA-1301765), and the State of Delaware. We thank Drs. Jeffrey Caplan and Sylvain Le Marchand for their expert assistance in confocal imaging and image analysis. We thank Sanofi/Genzyme for generously providing HA.

