## Supplemental Data 1 for "TGFβR Inhibition Represses TGF-β1 Initiated Keratin-7 Expression in Human Salivary Gland Progenitor Cells"

Dr. Eric W. Fowler, Prof. X. Jia
Department of Materials Science and Engineering
University of Delaware
Newark, Delaware, 19716, USA

Emmett V. Venrooy
Department of Biological Sciences
University of Delaware
Newark, Delaware, 19716, USA

M.D. Robert L. Witt
Helen F. Graham Cancer Center and Research Institute
Christiana Care Health Systems
Newark, Delaware, 19713, USA

***Simultaneous RNA, DNA and Protein Isolation from 3D Cultures***: On days 1, 3, 7, and 14 of culture, hS/PC laden HA hydrogel constructs were frozen with liquid nitrogen and stored at -80 °C until extraction was performed. Extractions were conducted by crushing the frozen hydrogel constructs with polypropylene Pellet Pestles™ (12-141-367; Fisher Scientific), followed by treatment with 750 μL TRIzol™ reagent. After complete homogenization in TRIzol™, the samples were centrifuged at 4 °C for 5 min at 15,000 × g. The resulting pellet with insoluble cross-linked HA hydrogel was temporarily stored at 4 °C for subsequent isolation of DNA and protein. The TRIzol™ supernatant was collected, and chloroform (150 μL) was added before incubating samples for 3 min at room temperature. The supernatant was then centrifuged at 4 °C for 20 min at 15,000 × g to yield a phase-separated solution. The lower phase was collected for DNA and protein isolation and temporarily stored at 4 °C. The upper aqueous phase was collected, and RNA purification was conducted using a RNA Clean & Concentrator™-5 Kit (R1013; Zymo Research, Irvine, CA) according to the manufacturer's protocol. After column elution, RNA quantification and purity were assessed using a NanoDrop™ 2000 Spectrophotometer (Nanodrop Technologies, Wilmington, DE). This technique yielded high purity RNA with absorbance ratios 260/280 nm >1.95 and 260/230 >1.8.

To continue the DNA isolation, the insoluble fraction generated from the initial TRIzol™ homogenized HA hydrogel was recombined with the corresponding lower TRIzol™ phase and briefly vortexed to permit dissolution. Ethanol (100%, 2.5 mL) was then added, and samples were mixed by inversion, incubated at room temperature for 3 min, then centrifuged at 2000 × g at 4 °C for 5 min. The phenol-ethanol supernatant was collected and stored at 4 °C until further protein isolation described below. The resulting pellet was incubated with 750 μL of 0.1 ᴍ sodium citrate containing 10% (v/v) ethanol and mixed by gentle inversion for 30 min at room temperature. The DNA pellet was centrifuged at 4 °C for 5 min at 2,000 × g, and the procedure was conducted for a total of three treatments of the solution containing 0.1 ᴍ sodium citrate/10% (v/v) ethanol. The DNA pellet was dissolved in pH 9.5 TE buffer and stored at -80 °C until further characterization.

To continue the protein isolation, the phenol-ethanol supernatant was combined with isopropanol (1.5 mL) and incubated for 1 h at room temperature before an extended incubation was conducted for 16 h at 4 °C. Samples were then centrifuged at 4 °C for 1 h at 15,000 × g. The resulting protein pellet was then incubated at room temperature for 20 min with 0.3 ᴍ guanidine hydrochloride (GdnHCl) in 95% (v/v) ethanol (1.8 mL) before centrifuging at 4 °C for 20 min at 15,000 × g. Two additional GdnHCl washes were performed before washing with 100% ethanol. The protein pellets were next air-dried before dissolving in a solution of Tris HCl (100 mᴍ, pH 8.0), prepared with 4 ᴍ urea, 5% (w/v) sodium dodecyl sulfate, 140 mᴍ NaCl, 20 mᴍ ethylenediaminetetraacetic acid, and 10% (v/v) glycerol, then stored at -20 °C until further characterization.

***dsDNA Quantification***: DNA solutions obtained via TRIzol™ extraction were centrifuged at 12,000 × g for 10 min at 4 °C. The resulting supernatant was analyzed using a Quant-iT™ PicoGreen™ dsDNA Assay Kit following the manufacturer's protocol. Fluorescent measurements were conducted using a SpectraMax® i3x Multi-Mode Microplate Reader (Molecular Devices, San Jose, CA) with a 485 nm excitation and 520 nm emission wavelength.

***Western Blotting***: The protein concentration of cell lysates, obtained by TRIzol™ extraction, were quantified using a Micro BCA™ Protein Assay Kit (23235; Thermo Scientific™, Waltham, MA), according to the manufacturer's procedure, and measurements were performed on a SpectraMax® i3x Multi-Mode Microplate Reader at 562 nm. The cell lysate was then combined with 3 × Blue Loading Buffer (56036; Cell Signaling Technology®, Danvers, MA; 187.5 mм Tris-HCl, pH 6.8, 6% (w/v) SDS, 30% glycerol and 0.03% (w/v) bromophenol blue) and prepared with 0.2 M DTT (7016L; Cell Signaling Technology®). Next, samples were incubated for 5 min at room temperature before heating at 95 °C for 10 min and vortexed for 30 s before centrifuging at 15,000 × g for 2 min at room temperature. SDS-PAGE was conducted using 4–20% Mini-PROTEAN® TGX Stain-Free™ Protein Gels (4561093; Bio-Rad Laboratories, Hercules, CA) incubated with Tris-Glycine-SDS buffer (1610732; Bio-Rad Laboratories). Sample lysate was loaded at 10 µg per lane and applied with 40 volts for 30 min followed by 300 volts for 20 min at 4 °C. Samples were transferred to nitrocellulose membranes (12369P2; Cell Signaling Technology) at 4 °C for 2 h using 70 volts. For protein lane normalization, the nitrocellulose membranes were treated with Revert™ 700 Total Protein Stain (926-11010; LI-COR® Biosciences, Lincoln, NE) according to the manufacturer's protocol and imaged using an iBright™ FL1500 Imaging System (Invitrogen™, Carlsbad, CA). Membranes were blocked with 5% (w/v) skim milk (sm/TBST) in tris-buffered saline containing 0.1% Tween 20 (TBST, 9997; Cell Signaling Technology®) for 2 h at room temperature. Primary antibodies (Table S3) were then diluted in sm/TBST and incubated for 16 h at 4 °C. Membranes were next washed three times with TBST for 5 min at room temperature. Alkaline phosphatase-linked secondary antibodies (Cell Signaling Technology®, Table S3) were diluted 1/20,000 in sm/TBST and incubated for 1 h at room temperature before washing with TBST three times at room temperature. Chemifluorescence signal was introduced using Cytiva Amersham™ ECF™ substrate (RPN5785; MilliporeSigma) and visualized using an iBright™ FL1500 Imaging System. Densitometry analysis was conducted with ImageJ using the analyze gels feature, in agreement with the procedure outlined in the ImageJ documentation. (https://imagej.nih.gov/ij/docs/menus/analyze.html#gels)

***qPCR Assay*:** RNA was reverse transcribed using the QuantiTect® Reverse Transcription Kit (205314; QIAGEN, Hilden, Germany) following the manufacturer's protocol. Sequence-specific amplification and detection was performed on an ABI 7300 real-time PCR system (Applied Biosystems®, Foster City, CA), with a thermal cycling profile of 1 cycle at 95 °C for 10 min, followed by 40 cycles of 95 °C for 15 s and 60 °C for 1 min. Twenty μL PCR reactions were prepared in 96 well format by combining Power SYBR™ green PCR master mix (Applied Biosystems®), cDNA, and target specific primers. Primer designs were primarily sourced from PrimerBank,[1] (https://pga.mgh.harvard.edu/primerbank/) and the complete primer sequences are available in Table S1. Primers were synthesized by Integrated DNA Technologies (Coralville, IA) and utilized after confirming to amplify a single target on agarose gels with an amplicon size corresponding to that determined from Primer-BLAST.[2] Glyceraldehyde 3-phosphate dehydrogenase (GAPDH) was used as a reference gene, and inter-run calibrations were included when gene expression was examined across multiple time points.[3] Cycle threshold values were generated using 7300 System SDS RQ Study software version 1.4 (Applied Biosystems), and mRNA levels were resolved using Qbase+ software version 3.2 (Biogazelle, Zwijnaarde, Belgium). Three biological replicates are reported from three technical replicates measured in duplicate.

**Table S1.** Primers used for qPCR assays.

| Target | Forward Primer (5'-3') | Reverse Primer (5'-3') |
|---|---|---|
| *GAPDH* | CAGCCTCAAGATCATCAGCA | TGTGGTCATGAGTCCTTCCA |
| *KRT5* | CGTGCCGCAGTTCTATATTCT | ACTTTGGGTTCTCGTGTCAG |
| *KRT14* | CACAGATCCCACTGGAAGAT | GATAATGAAGCTGTATTGATTGCC |
| *TFCP2L1* | GCCGCCTGCTTCCTGTTC | CTGCCCACCACTGCTCAAAG |
| *AMY1A* | CTCGGCACAGTTATTCGCAAGTGG | ACAGCCTAGCATCCCAGAAGGT |
| *SLC12A2* | TAAAGGAGTCGTGAAGTTTGGC | CTTGACCCACAATCCATGACA |
| *KRT7* | AAGAACCAGCGTGCCAAGT | TCCAGCTCCTCCTGCTTG |
| *KRT18* | GTTGACCGTGGAGGTAGA | GACCCAGCTCGTCATATTGGG |
| *KRT19* | CTGCCTCCAAGGTCCTCT | CCCATCCCTCTACCCAGAAG |
| *KRT7-AS* | TCCAACGCCTATGTTCCAGTTC | ACATTGTGCCACGGACATCTTG |
| *TP53* | CAGCACATGACGGAGGTTGT | TCATCCAAATACTCCACACGC |
| *TGFB1* | GCAGAAGTTGGCATGGTAGC | CCCTGGACACCAACTATTGC |
| *GDF15* | GACCCTCAGAGTTGCACTCC | GCCTGGTTAGCAGGTCCTC |

| | | |
|---|---|---|
| *YAP1* | ACCCACAGCTAGCATCTTCG | TGGCTTGTTCCCATCCATCAG |
| *CTGF (CCN2)* | AGGAGTGGGTGTGTGACGA | CCAGGCAGTTGGCTCTAATC |
| *CYR61 (CCN1)* | CCTTGTGGACAGCCAGTGTA | ACTTGGGCCGGTATTTCTTC |
| *CCND1* | TGGAGCCCGTGAAAAAGAGC | TCTCCTTCATCTTAGAGGCCAC |
| *JUNB* | ACGACTCATACACAGCTACGG | GCTCGGTTTCAGGAGTTTGTAGT |
| *SMAD3* | TGGACGCAGGTTCTCCAAAC | CCGGCTCGCAGTAGGTAAC |
| *MYC* | CGGAACTCTTGTGCGTAAGG | TCATAGGTGATTGCTCAGGACAT |
| *EREG* | GTGATTCCATCATGTATCCCAGG | GCCATTCATGTCAGAGCTACACT |
| *AREG* | GTGGTGCTGTCGCTCTTGATA | CCCCAGAAAATGGTTCACGCT |
| *HBEGF* | ATCGTGGGGCTTCTCATGTTT | TTAGTCATGCCCAACTTCACTTT |
| *TGFA* | AGGTCCGAAAACACTGTGAGT | AGCAAGCGGTTCTTCCCTTC |
| *ADAM10* | ATGGGAGGTCAGTATGGGAATC | ACTGCTCTTTTGGCACGCT |
| *ADAM17* | GTGGATGGTAAAAACGAAAGCG | GGCTAGAACCCTAGAGTCAGG |
| *TIMP3* | CATGTGCAGTACATCCATACGG | CATCATAGACGCGACCTGTCA |
| *TIMP1* | TTTCTTGGTTCCCCAGAATG | CAGAGCTGCAGAGCAACAAG |
| *MMP1* | GGGAGATCATCGGGACAACTC | GGGCCTGGTTGAAAAGCAT |
| *FN* | ACCTACGGATGACTCGTGCTTTGA | CAAAGCCTAAGCACTGGCACAACA |
| *LAMA1* | GTGATGGCAACAGCGCAAA | GACCCAGTGATATTCTCTCCCA |
| *ITGA5* | GCCTGTGGAGTACAAGTCCTT | AATTCGGGTGAAGTTATCTGTGG |
| *ITGAV* | AATCTTCCAATTGAGGATATCAC | AAAACAGCCAGTAGCAACAAT |
| *IL6* | ACTCACCTCTTCAGAACGAATTG | CCATCTTTGGAAGGTTCAGGTTG |
| *CXCL8* | ACATACTCCAAACCTTTCACCC | CAACCCTCTGCACCCAGTTTTC |
| *IL10* | GGTTGCCAAGCCTTGTCTGA | AGGGAGTTCACATGCGCCT |
| *IL1B* | GCTGATGGCCCTAAACAGATGA | TTGCTGTAGTGGTGGTCGGAGAT |
| *CDKN1A (P21)* | TGTCCGTCAGAACCCATGC | AAAGTCGAAGTTCCATCGCTC |
| *CDKN2A (P16)* | GATCCAGGTGGGTAGAAGGTC | CCCCTGCAAACTTCGTCCT |
| *SERPINE1* | ACCGCAACGTGGTTTTCTCA | TTGAATCCCATAGCTGCTTGAAT |
| *JUNB* | ACGACTCATACACAGCTACGG | GCTCGGTTTCAGGAGTTTGTAGT |
| *IGF1* | GCTCTTCAGTTCGTGTGTGGA | GCCTCCTTAGATCACAGCTCC |
| *IGF2* | GTGGCATCGTTGAGGAGT | CACGTCCCTCTCGGACTT |
| *IGF1R* | TCGACATCCGCAACGACTATC | CCAGGGCGTAGTTGTAGAAGA |

**Table S2.** Antibodies and conditions used for 3D immunocytochemistry.

| Target | Vendor | Clone/<br>Item No./RRID | Host | Perm<br>(w/v) | Dilution<br>(v/v) | Secondary Antibody |
|---|---|---|---|---|---|---|
| YAP1 | Santa Cruz Biotechnology | (63.7)/<br>sc-101199<br>RRID:AB_1131430 | Ms | 0.2%<br>Triton | 1/50 | Alexa Fluor® 647 AffiniPure Fab Fragment Goat Anti-Ms IgG (H+L) AB_2338931 |
| SMAD 2/3 | Cell Signaling Technology | (D7G7) XP®/<br>8685S<br>RRID:AB_10889933 | Rb | 0.2%<br>Triton | 1/100 | Alexa Fluor® 488 AffiniPure Fab Fragment Goat Anti-Rb IgG (H+L)<br>AB_2338058 |
| Keratin 5 | BioLegend | 905501/<br>RRID:AB_2565050 | Rb | 0.2%<br>Saponin | 1/100 | Alexa Fluor® 647 AffiniPure Fab Fragment Goat Anti-Rb IgG (H+L)<br>AB_2338084 |
| β-catenin | Santa Cruz Biotechnology | (E-5)/<br>sc-7963<br>RRID:AB_626807 | Ms | 0.2%<br>Saponin | 1/25 | Alexa Fluor® 488 AffiniPure Fab Fragment Goat Anti-Ms IgG (H+L)<br>AB_2338869 |
| α-Amylase | MilliporeSigma | A8273/<br>RRID:AB_258380 | Rb | 0.05%<br>Saponin | 1/100 | Alexa Fluor® 488 AffiniPure Fab Fragment Goat Anti-Rb IgG (H+L)<br>AB_2338058 |

**Table S3.** Antibodies and conditions used for western blotting.

| Target | Vendor | Clone/<br>Item No./RRID | Host | Dilution<br>(v/v) | Secondary Antibody |
|---|---|---|---|---|---|
| Keratin 5 | BioLegend | 905501<br>RRID:AB_2565050 | Rb | 1/20,000 | Anti-rabbit IgG, AP-linked Antibody #7054[i]<br>RRID:AB_2099235 |
| Fibronectin | MilliporeSigma | F3648<br>RRID:AB_476976 | Rb | 1/1000 | Anti-rabbit IgG, AP-linked Antibody #7054[i]<br>RRID:AB_2099235 |
| Keratin 7 | MilliporeSigma | OV-TL 12/30/<br>MAB3554<br>RRID:AB_94924 | Ms | 1/1000 | Anti-mouse IgG, AP-linked Antibody #7056[i]<br>RRID:AB_330921 |
| Keratin 14 | Abcam | LL002/<br>Ab7800<br>RRID:AB_306091 | Ms | 1/1000 | Anti-mouse IgG, AP-linked Antibody #7056[i]<br>RRID:AB_330921 |

(i) Cell Signaling Technology

**Table S4.** Antibodies and conditions used for 2D immunocytochemistry.

| Target | Vendor | Clone/<br>Item No./RRID | Host | Perm<br>(w/v) | Dilution<br>(v/v) | Secondary Antibody |
|---|---|---|---|---|---|---|
| YAP1 | Santa Cruz Biotechnology | (63.7)/<br>sc-101199<br>RRID:AB_1131430 | Ms | 0.2%<br>Triton | 1/50 | Alexa Fluor® 647 Goat anti-Ms<br>RRID:AB_2535805 |
| SMAD 2/3 | Cell Signaling Technology | (D7G7) XP®/<br>8685S<br>RRID:AB_10889933 | Rb | 0.2%<br>Triton | 1/100 | Alexa Fluor® 488 Goat anti-Rb<br>RRID:AB_2576217 |

| | | | | | | |
|---|---|---|---|---|---|---|
| Keratin 5 | BioLegend | 905501<br>RRID:AB_2565050 | Rb | *0.05%<br>Saponin | 1/100 | Alexa Fluor® 647<br>Goat anti-Rb[ii]<br>RRID:AB_2535813 |
| Keratin 7 | MilliporeSigma | OV-TL 12/30/<br>MAB3554<br>RRID:AB_94924 | Ms | 0.05%<br>Saponin | 1/50 | Alexa Fluor® 488<br>Goat anti-Ms<br>RRID:AB_2534088 |
| Ki-67 | Invitrogen | 7B11/ 334711/<br>RRID: AB_2533122 | Ms | 0.2% Triton | 1/450 | *NA* (FITC conjugated) |

(ii) K5/YAP1 ICC was performed with 0.2 % Triton (w/v) and Alexa Fluor® 488 goat anti-Rb

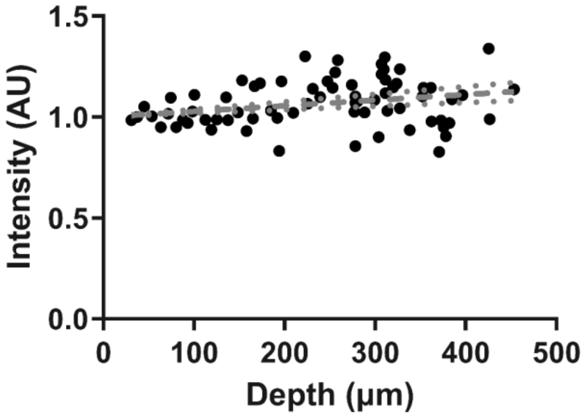


**Figure S1.** Alexa Fluor® 647 labeled reference microsphere standards were suspended in HA hydrogels, and z-stack images were captured with fluorescent microscopy. The mean intensity of individual microspheres (black circles) was determined using Imaris 3D-4D imaging software. Linear regression of microsphere normalized mean intensity calculated using GraphPad Prism 9 (GraphPad Software, San Diego, CA) is shown in gray; dotted red circles represent 95% confidence intervals.

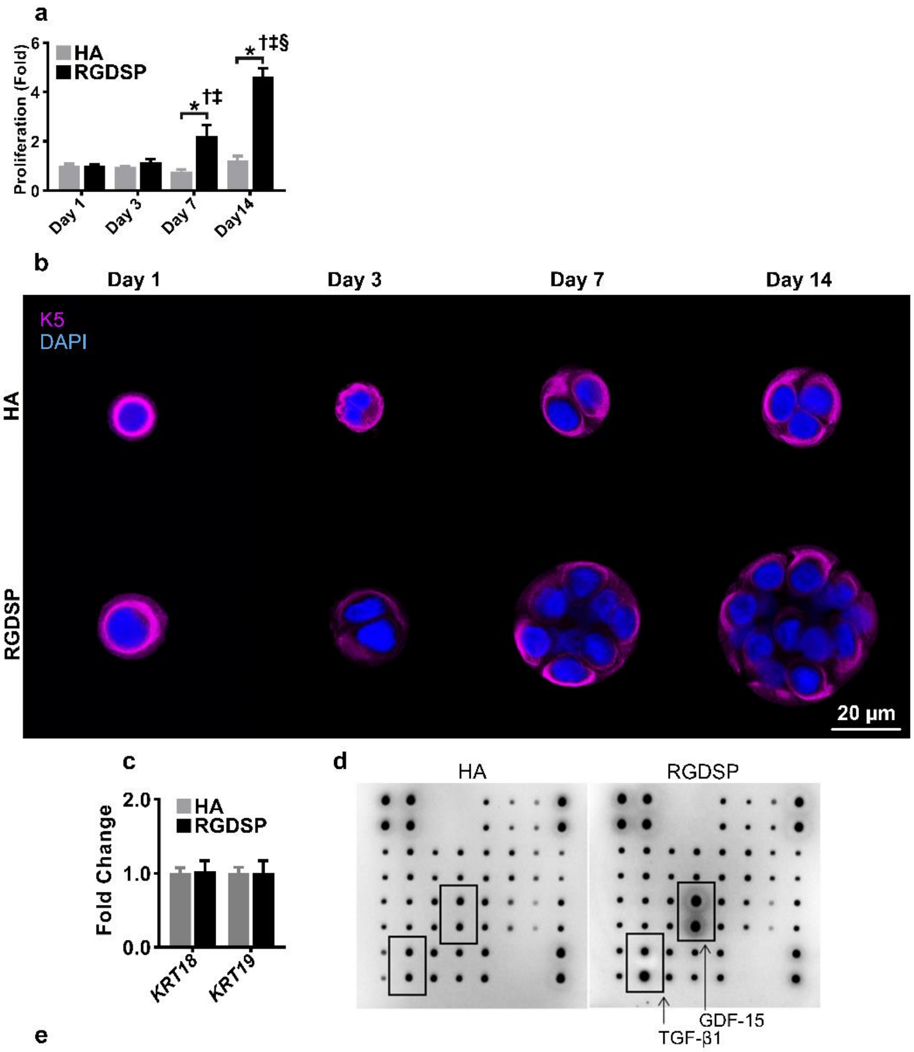


e

Each antibody is spotted in duplicate vertically

| | A | B | C | D | E | F | G | H |
|---|---|---|---|---|---|---|---|---|
| | POS | POS | NEG | NEG | Activin A | Activin R1A | BAMBI | beta-IG-H3 |
| | BMP2 | BMP4 | BMP6 | BMP7 | BMP8 | BMP9 | BMP15 | BMPRIA |
| | BMPRII | Follistatin | GDF11 | GDF15 | LeftyA | NOGGIN | SMAD3 | SMAD4 |
| | TGFa | TGFb1 | TGFb2 | TGFbRII | TGFbRIII | NEG | NEG | POS |

**Figure S2.** (**a**) hS/PCs were cultured in HA, and RGDSP hydrogels and DAPI stained nuclei were recorded with fluorescent microscopy before cell density quantification was performed with Imaris 3D Software; two-way ANOVA was performed followed by Tukey's multiple comparison test. * indicates $p < 0.05$ between HA and RGDSP at the same time point. †, ‡, § indicates $p < 0.05$ from day 1, 3, and day 7 measurements of the same data set, respectively. (**b**) hS/PCs were encapsulated in HA and RGDSP hydrogels, and K5 was visualized with fluorescent microscopy on days 1, 3, 7, and 14 of culture. (**c**) hS/PCs were cultured in HA and RGDSP hydrogels for 14 days, and qPCR was performed to determine *KRT18* and *KRT19* expression; Student's t-test, non-significant $p >$

0.05. (**d**) TGF-β Superfamily immunoblot array detailing elevated TGF-β1 and GDF-15 secreted by RGDSP cultures. (**e**) TGF-β Superfamily immunoblot array template. Error bars represent SEM in all cases.

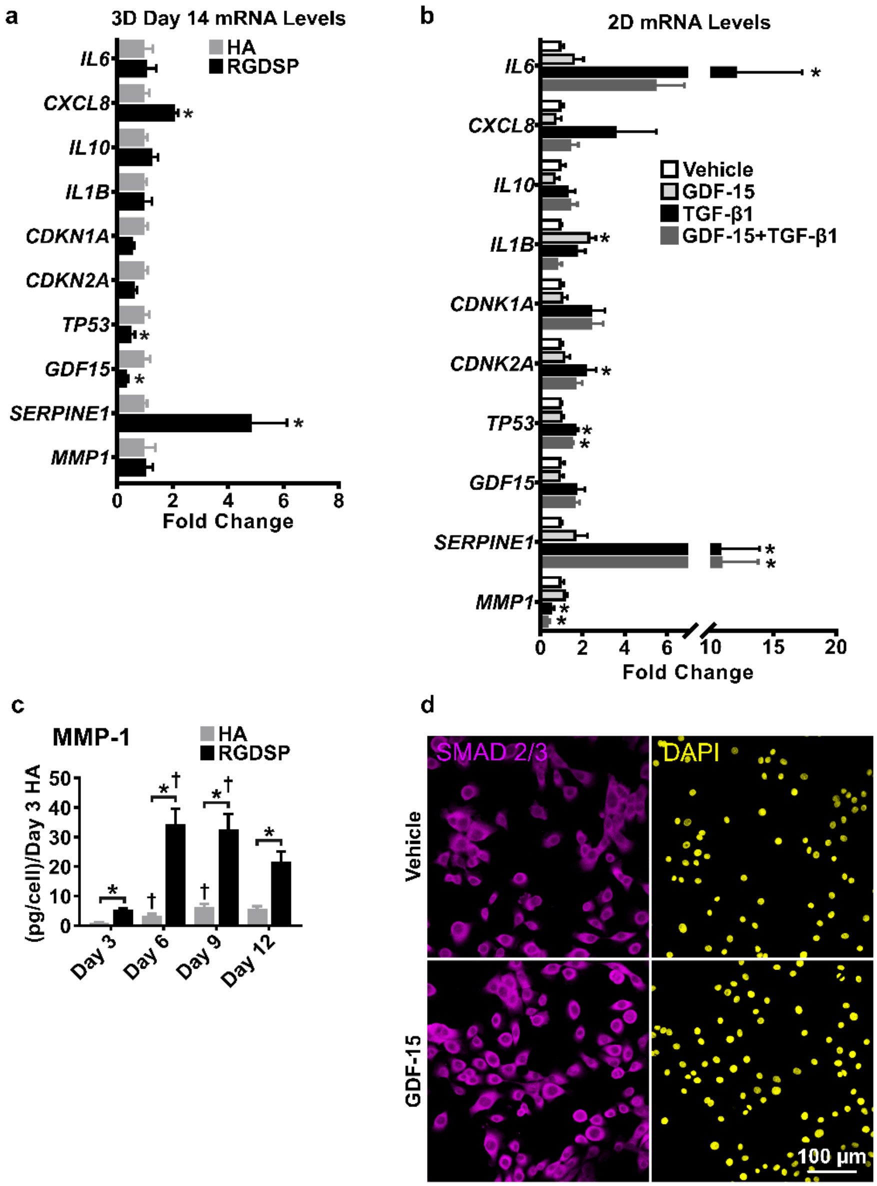


**Figure S3.** (**a**) hS/PCs were cultured for 14 days in HA, and RGDSP constructs and the expression of SASP associated genes were assessed by qPCR; Student's t-test, * indicates $p < 0.05$. (**b**) 2D cultured hS/PCs were treated with TGF-β1, GDF-15, and GDF-15+TGF-β1 for 48 h before assessing expression of SASP associated genes; one way-ANOVA was performed followed by a Dunnett's test. * indicates $p < 0.05$. (**c**) Secreted MMP-1 in HA and RGDSP cultures was determined by ELISA; two-way ANOVA was performed, followed by Tukey's multiple comparison test. * indicates $p < 0.05$ between HA and RGDSP at the same time point while. †, ‡, § indicates $p < 0.05$ from day 3, 6, and day 9 measurements of the same data set, respectively. (**d**) hS/PCs were treated with GDF-15 for 48 h, and the expression of SMAD 2/3 was visualized with fluorescent microscopy. Error bars represent SEM in all cases.

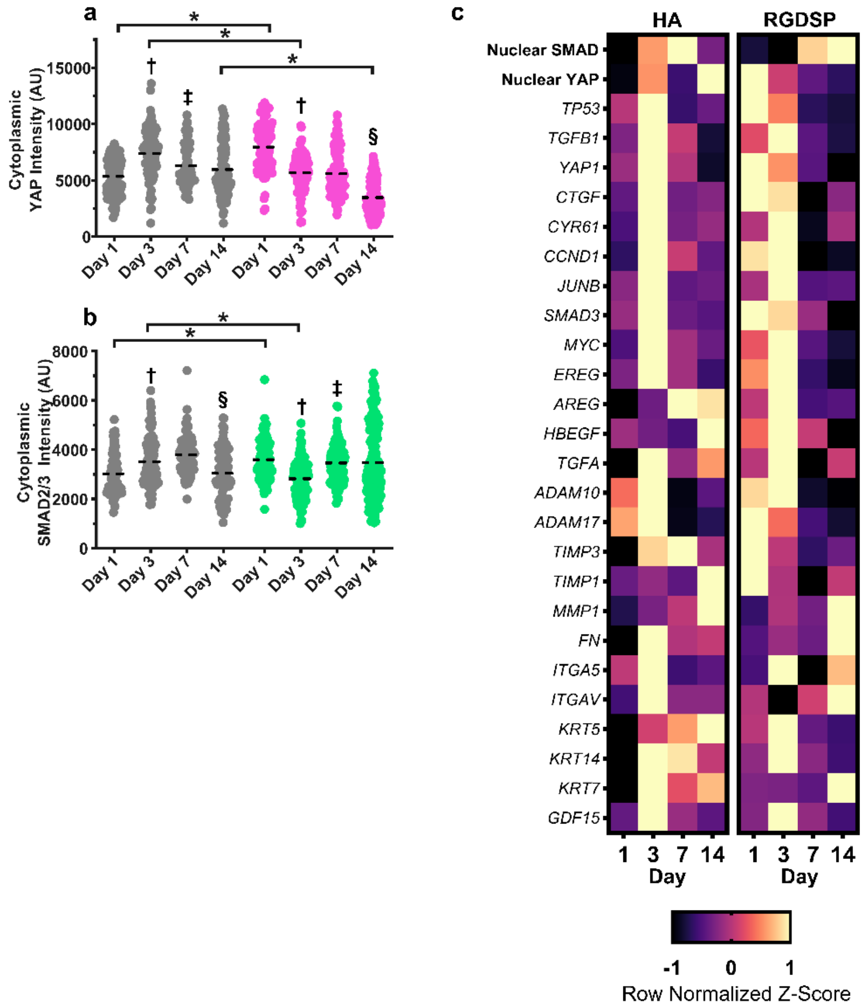


**Figure S4.** (**a-b**) hS/PCs were cultured in HA and RGDSP, and the cytoplasmic expression of YAP (a) and SMAD 2/3 (b) was resolved on days 1, 3, 7, and 14 using ICC. Filled circles represent individual cells or multicellular structures as spheroids develop, and the dashed black line indicates the mean value of each data set. Two-way ANOVA was performed, followed by Tukey's multiple comparison test. * indicates $p < 0.05$ between HA and RGDSP at the same time point. †, ‡, § indicates $p < 0.05$ from day 1, 3, and day 7 measurements of the same data set, respectively. (**c**) The expression of TGF-β and YAP target genes was assessed by qPCR on days 1, 3, 7, and 14 of HA and RGDSP cultures. Z-scores are reported as row normalized expression to aid comparisons with nuclear SMAD and YAP intensities of HA and RGDSP cultures. Error bars represent SEM in all cases.

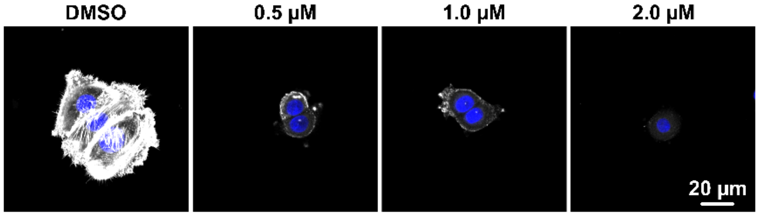


**Figure S5**. hS/PC were cultured with VERT (0.5, 1.0, and 2.0 μM) and a DMSO for 24 h. ICC was performed to investigate the expression of F-actin (white). Cell nuclei were labeled with DAPI (blue).

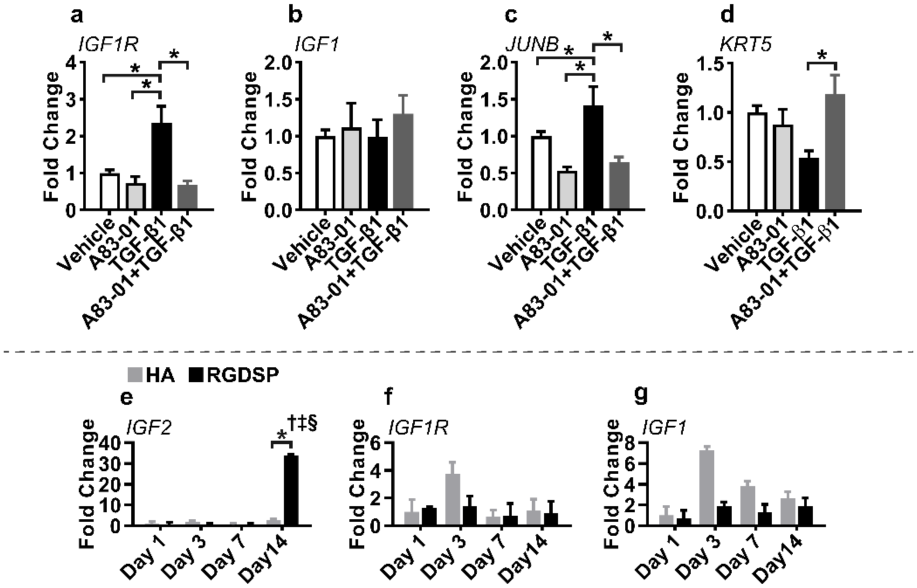


**Figure S6. (a-d)** hS/PC were cultured with TGF-β1, A83-01, or in combination for 48 h before the expression of *IGF1R* (a) *IGF1* (b) *JUNB* (c), and *KRT5* (d) was assessed with qPCR; One way-ANOVA was performed followed by Tukey's multiple comparison test. * indicates $p < 0.05$. (**e-g**) hS/PCs were cultured in HA, and RGDSP hydrogels and *IGF2* (e), *IGF1R* (f), and *IGF1 (g)* expression was investigated with qPCR on days 1, 3, 7, and 14. Two-way ANOVA was performed, followed by Tukey's multiple comparison test. * indicates $p < 0.05$ between HA and RGDSP at the same time point. $^{†, ‡, §}$ indicates $p < 0.05$ from day 1, 3, and day 7 measurements of the same data set, respectively.
